# Donor Macrophages Modulate Rejection after Heart Transplantation

**DOI:** 10.1101/2021.09.17.459296

**Authors:** BJ Kopecky, H Dun, JM Amrute, CY Lin, AL Bredemeyer, Y Terada, PO Bayguinov, AL Koenig, CC Frye, JAJ Fitzpatrick, D Kreisel, KJ Lavine

## Abstract

**Background:** Cellular rejection after heart transplantation imparts significant morbidity and mortality. Current immunosuppressive strategies are imperfect, target recipient T-cells, and have a multitude of adverse effects. The innate immune response plays an essential role in the recruitment and activation of T-cells. Targeting the donor innate immune response would represent the earliest interventional opportunity within the immune response cascade. There is limited knowledge regarding donor immune cell types and functions in the setting of cardiac transplantation and no current therapeutics exist for targeting these cell populations.

**Methods:** Using genetic lineage tracing, cell ablation, and conditional gene deletion, we examined donor mononuclear phagocyte diversity and function during acute cellular rejection of transplanted hearts in mice. We performed single cell RNA sequencing on donor and recipient macrophages, dendritic cells, and monocytes at multiple timepoints after transplantation. Based on our single cell RNA sequencing data, we evaluated the functional relevance of donor CCR2^+^ and CCR2^-^ macrophages using selective cell ablation strategies in donor grafts prior to transplant. Finally, we perform functional validation of our single cell-derived hypothesis that donor macrophages signal through MYD88 to facilitate cellular rejection.

**Results:** Donor macrophages persisted in the transplanted heart and co-existed with recipient monocyte-derived macrophages. Single-cell RNA sequencing identified donor CCR2^+^ and CCR2^-^ macrophage populations and revealed remarkable diversity amongst recipient monocytes, macrophages, and dendritic cells. Temporal analysis demonstrated that donor CCR2^+^ and CCR2^-^ macrophages were transcriptionally distinct, underwent significant morphologic changes, and displayed unique activation signatures after transplantation. While selective depletion of donor CCR2^-^ macrophages reduced allograft survival, depletion of donor CCR2^+^ macrophages prolonged allograft survival. Pathway analysis revealed that donor CCR2^+^ macrophages were being activated through MYD88/NF-ĸβ signaling. Deletion of MYD88 in donor macrophages resulted in reduced antigen presenting cell recruitment, decreased emergence of allograft reactive T-cells, and extended allograft survival.

**Conclusions:** Distinct populations of donor and recipient macrophages co-exist within the transplanted heart. Donor CCR2^+^ macrophages are key mediators of allograft rejection and inhibition of MYD88 signaling in donor macrophages is sufficient to suppress rejection and extend allograft survival. This highlights the therapeutic potential of donor heart-based interventions.

## Introduction

Heart transplantation is the definitive treatment for end-stage heart failure with over 3000 transplants performed annually in the United States [1, 2]. Given the complexities of donor-recipient matching and imperfect immunosuppressive regimens, approximately 40% of transplanted patients suffer rejection within their first year after transplant leading to adverse short and long-term outcomes [3, 4]. Current strategies target recipient T-cells and impose harmful consequences of systemic immunosuppression including infection and malignancy [5].

T-cell activation by antigen presenting cells, including macrophages and dendritic cells, is a requisite step for allograft rejection [6–12]. Limiting the emergence and proliferation of alloreactive T-cells by targeting macrophages and/or dendritic cells may mitigate the incidence of allograft rejection while avoiding adverse outcomes associated with systemic T-cell depletion or inhibition. While recipient-derived macrophages and dendritic cells have an established role in heart transplant rejection [13–18], the precise role of donor-derived immune cells remain largely unexplored.

Donor hearts contains abundant populations of resident macrophages and fewer dendritic cells [19]. Targeting cardiac graft-resident macrophage activation represents an attractive approach to reduce rejection while eschewing systemic immunosuppression [15]. Interestingly, early activation of macrophages within the allograft can result in a prolonged activated inflammatory state that persists in the absence of antigen or other stimuli, a property termed “trained immunity” [20]. This suggests that early activation of donor macrophages may have long-term functional effects on allograft outcomes that are independent of ongoing immunosuppression.

We have previously reported that mouse and human hearts contain at least two subsets of resident macrophages with divergent origins and functions that can be distinguished based on the expression of C-C chemokine receptor 2 (CCR2). CCR2^+^ macrophages are derived from adult hematopoietic progenitors, are replenished through ongoing monocyte recruitment, and orchestrate inflammatory responses including monocyte and neutrophil infiltration. CCR2^-^ macrophages are derived from embryonic hematopoietic progenitors, are maintained independent of monocyte input, suppress inflammation, and promote tissue repair [7, 19, 21–24]. The role of donor CCR2^+^ and CCR2^-^ macrophages in the context of heart transplantation rejection remains to be elucidated [25].

In the present study, we utilize a murine heart transplantation model to precisely interrogate the roles of donor macrophages during acute cellular rejection [26, 27]. By combining genetic lineage tracing with single cell RNA sequencing, we examine transcriptional signatures of donor and recipient macrophages and identify putative signaling mechanisms underlying donor CCR2^+^ and CCR2^-^ macrophage activation. We use genetic lineage tracing to demonstrate that donor CCR2^+^ and CCR2^-^ macrophages persist after heart transplantation. Targeted cell depletion strategies show that reduction of donor CCR2^+^ macrophages result in prolonged allograft survival with reduced inflammation whereas depletion of donor CCR2^-^ macrophages conversely leads to rapid rejection with increased inflammation. We show that donor CCR2^+^ macrophages signal in part through MYD88 and inhibition of this signaling either in the donor graft or donor macrophages suppresses alloantigen-specific T-cell reactivity resulting in prolongation of allograft survival. Together, these findings establish donor CCR2^+^ macrophages as a viable therapeutic target to ameliorate or prevent heart transplant rejection.

## Materials and Methods

### Animal Models

Mice were bred and maintained at the Washington University School of Medicine and all experimental procedures were performed in accordance with the animal use oversight committee. Mouse strains utilized included CD169^DTR/+^ [28], CCR2^DTR/+^ [29], MyD88^f/f^ [30], LysM^Cre/+^ [31], CSF1r^ertCre/+^ [32], MyD88^-/-^ [33], CX3CR1^GFP/+^ CCR2^RFP/+^ [34, 35]. All donor mice were on the C57BL/6 (B6) background and genotyped according to established protocols [36]. Recipient mice were age- and gender-matched BALB/c mice between 6-8 weeks of age. Equal numbers of male and female mice were included in all experiments. Diphtheria toxin receptor (DTR) and control mice were given 200 ng intraperitoneal injections of diphtheria toxin (Sigma Cat #D0564) on three consecutive days prior to transplant. Tamoxifen food pellets (500 mg/kg diet, Envigo Teklad Diets 500 TD130857) were provided for two weeks prior to transplantation.

### Heterotopic Heart Transplantation

Heart grafts were harvested from donor mice and transplanted heterotopically into the abdomen of recipients following 1 hour of cold (4°C) ischemia, as previously described [26]. For low-dose immunosuppression, 200 µg CTLA4-Ig (Bio X Cell Cat BE0099) was administered intraperitoneally on post-transplant days 0, 2, 4, and 6. For <u>high-dose</u> immunosuppression, 1.25 mg CTLA4-Ig was administered on post-transplant days 0, 4, and 14. After transplantation, allografts were palpated daily. Cessation of a palpable heartbeat, confirmed by visual inspection, indicated rejection of the cardiac allograft. Perioperative graft loss (within 72 hours) was excluded from the analysis.

### Histology

After ice cold PBS perfusion, atria of donor hearts were removed. Ventricles were placed in 4% PFA overnight at 4°C. Hearts were then rinsed with PBS x 3 and placed in PBS at 4°C overnight. Hearts were then placed in histology cassettes for paraffin embedding and dehydrated in 70% ethyl alcohol. Tissues were paraffin embedded and 4 µm sections were cut and stained with H&E. Images were obtained with EVOS imaging system model FLc or an AxioScan Z1 (Carl Zeiss, Jena, Germany). After imaging, slides were scored in a blinded fashion by a trained cardiac pathologist based on the 1990 and 2004 ISHLT cellular rejection guidelines [37].

### Immunofluorescence

After ice cold PBS perfusion, ventricles were placed in 4% PFA overnight at 4°C. Hearts were rinsed with PBS x 3 and infiltrated with 30% sucrose overnight at 4°C. Hearts were embedded in O.C.T. (Fisher HealthCare Tissue Plus O.C.T. Compound Cat 4585) and frozen at -80°C. 10 or 30 µm sections were obtained using a Leica Cryostat. Sections were then washed in TBS and stained in 10% FBS in TBS-T (0.05% Tween-20) blocking solution with primary antibodies: anti-GFP 1:2000 (Abcam Cat# ab13970), anti-RFP 1:1000 (Rockland p/n: 600-401-379), anti-mouse CD45 1:200 (BD Pharmingen Cat# 550539), anti-mouse CD68 1:200 clone FA-11 (Biolegend Cat# 137002), anti-Ki-67 1:200 (Invitrogen Cat# 14-5698-82) overnight at 4°C in a humidified environment. For TUNEL staining, prior to primary antibody staining, Click-iT Plus TUNEL Assay with Alexa Fluor 647 dye Cat# C10619 was used followed by standard antibody staining as above. After washing, appropriate secondary antibodies were added to blocking buffer and sections were stained for 60 minutes at room temperature protected from light (Alexa Fluor 488 goat anti-chicken (Cat # A11039), Alexa Fluor 555 goat anti-rabbit (Cat # A21428), Alexa Fluor 647 goat anti-rat (Cat# A21247)). DAPI mounting, anti-fade solution (Vectashield Cat #H-1200) was added immediately prior to placement of no. 1.5 coverslips. Images were obtained with Zeiss LSM 700 confocal microscope installed on an AxioImager.M2. Acquired images were processed using ZEN Blue and/or Black or Imaris V9.5 (Oxford Instruments). When quantifying cells, 10 µm sections were used. Entire heart sections were imaged with a 20X objective lens using both tile and z-stack features. Z-stacks were collapsed, and tiles stitched in ZEN. Cells were counted when the antibody of interest co-localized with DAPI. When possible, entire heart sections were counted. For 3D reconstruction, 30 µm sections were imaged with a single 40X z-stack. For surface area, volume, and projection quantification, at least two 40X fields in at least two non-consecutive sections were used for each heart. When possible, between 10-20 donor CCR2^-^ or CCR2^+^ macrophages were averaged for each heart. To assess cellular projections, reconstructed images were rotated and only distinct projections that measured at least 10 µm were considered. In all cases, genotype identification was blinded to the observer.

### ELISPOT

10 days after transplant, recipient BALB/c splenocytes were frozen using C.T.L.-Cryo ABC Media (ImmunoSpot Cat#CTLC-ABC). In brief, fresh spleens were pressed through a 40 μm cell strainer and rinsed with 1X C.T.L. wash (ImmunoSpot Cat# CTLW-010). 1:1 CTL-C and CTL-A/B were added. Up to 10 million splenocytes /mL were aliquoted into 1 mL cyrovials, which were subsequently stored in liquid nitrogen. Responder splenocytes were thawed using C.T.L. anti-aggregate wash (ImmunoSpot Cat# CTL-AA-005), T-cells were isolated (Miltenyi Pan T Cell isolation kit II (130-095-130)) and used for ELISPOT assay. Irradiated (20cGy) B6 (stimulator-allogeneic) and BALB/c (stimulator-syngeneic) splenocytes were added to responder T-cells in C.T.L. Test media (ImmunoSpot Cat# CTLT-010) at a ratio of 600,000 stimulators to 150,000 responder T-cells overnight at 37°C. IFN-γ spots were detected per C.T.L. ELISPOT protocol (ImmunoSpot Mouse IFN-γ Single-Color ELISPOT). Plates were analyzed and quality controlled by a blinded third-party using CTL-ImmunoSpot S6 University Analyzer. Positive (ConA) and negative controls were included. All samples were performed in technical triplicates and each experimental condition consisted of at least six independent samples.

### Flow Cytometry

Single cell suspensions were generated from ice cold PBS perfused hearts by finely mincing and digesting ventricles in DMEM with Collagenase IV (Sigma C5138) 4500 U/mL in DMEM, Hyaluronidase 1 (Sigma H3506) 2400 U/mL in DMEM, and DNAse I (Sigma D4527-40KU) 6000 U/mL in DMEM for 45 minutes at 37°C. To deactivate the enzymes, samples were washed with HBSS that was supplemented with 2% FBS and 0.2% BSA and filtered through 40 μm cell strainers. Red blood cell lysis was performed with ACK lysis buffer (Thermo Fisher Scientific) for five minutes at room temperature. Samples were washed with DMEM and resuspended in 100 μL of FACS buffer (DPBS with 2% FBS and 2 mM EDTA). Cells were stained with monoclonal antibodies at 4°C for 30 minutes in the dark. A complete list of antibodies is provided below. Samples were washed in FACS buffer and final resuspension was made in 300 μL FACS buffer. DAPI was used for identification of dead cells. Immune cells were first gated as CD45^+^ followed by standard doublet exclusion. For single cell RNA sequencing sorting, flow cytometric analysis and sorting were performed on a BD FACS ARIAIII platform. Monocytes, macrophages, and dendritic cells were first gated as CD11b^+^ and then all Ly6C^+^ and CD64^+^ cells were further assessed. Donor (GFP^+^) and recipient (GFP^-^) cells were sorted separately. Donor macrophages were identified as CD45^+^ CD11b^+^ Ly6c^-^ CD64^+^ GFP^+^ and then classified as alive (DAPI^-^) or dead (DAPI^+^). Spleens were pressed through 40 μM cell strainers and rinsed with DMEM. They were not further digested. Preparation was otherwise identical to heart single cell suspension/staining. Cell death flow cytometric analyses were performed on BD FACS Melody.

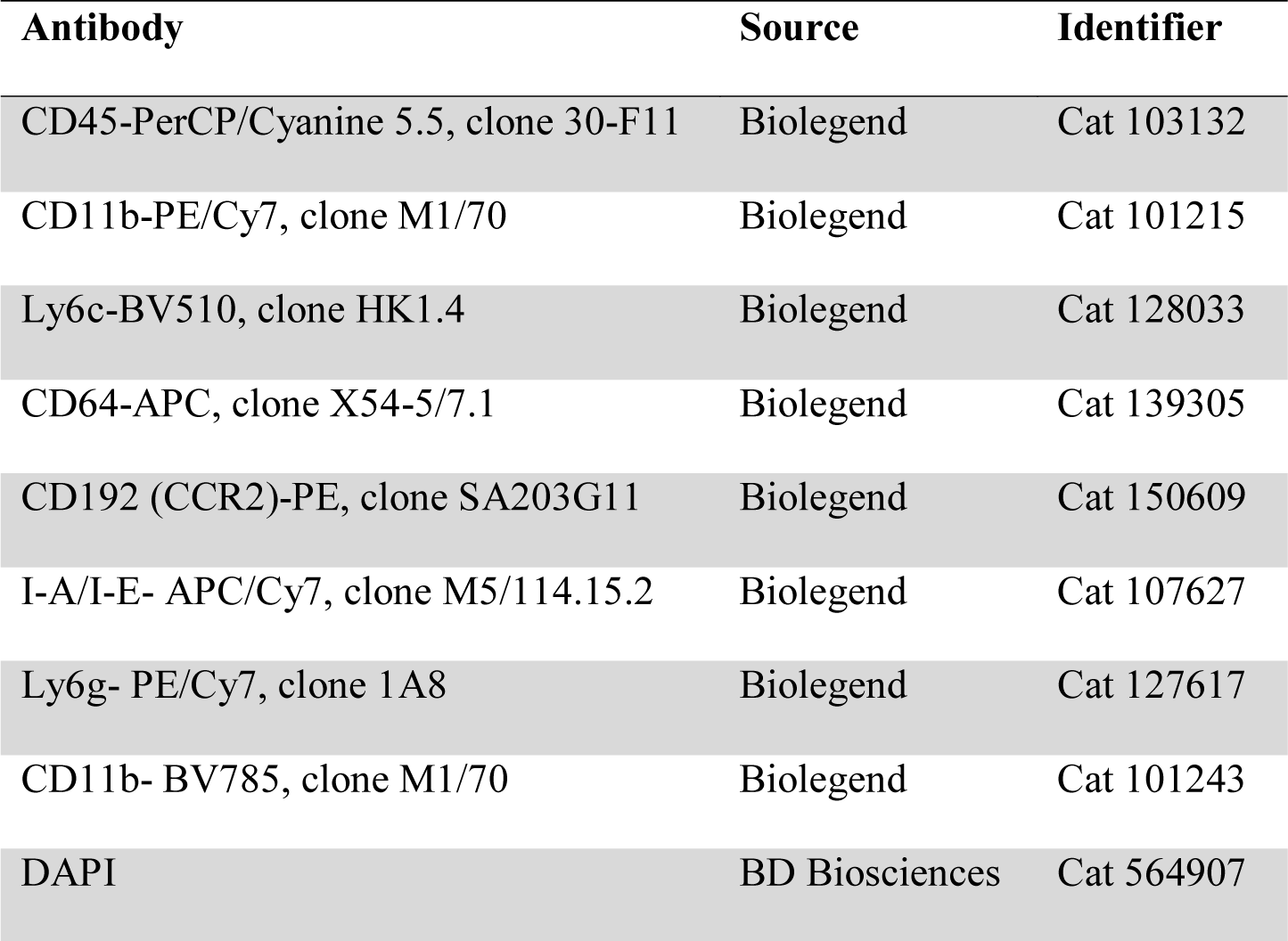

FACS sorted CD45^+^CD11b^+^Ly6G^-^CD64^+^ (Ly6c^+/-^) cells were processed and encapsulated with barcoded oligo-dT containing gel beads with the 10X Genomics Chromium controller. Library preparation was performed as per manufacturer recommended protocols at the McDonnel Genome Institute at Washington University. Single cell libraries were multiplexed into a single lane and were sequenced at a target read depth of 100,000 reads/cell using a NovaSeq sequencer (Illumina).

### Single Cell Analysis

Pre-processing: Sequencing alignment and de-multiplexing was performed using Cell Ranger from 10X Genomics to generate feature barcoded count matrices. All subsequent analysis was performed using the Seurat v4.0.0 package. The following quality control steps were performed to filter the count matrices: 1) genes expressed in fewer than 3 cells and cells expressing fewer than 200 genes were removed; 2) cells expressing > 5,000 genes and > 50,000 counts were discarded as these could be potential multiplet events; 3) cells with > 10% mitochondrial content were filtered out as these were deemed to be of low-quality. Normalization and variance-stabilization of raw counts was performed using SCTransform to find 3,000 variably expressed genes and percentage mitochondrial reads were regressed out. Principle component analysis was used to find nearest neighbors and a 2D UMAP embedding was used for visualization. Differential gene expression testing was performed using the FindAllMarkers function between conditions and across clusters. Statistically significant genes (adjusted p-value < 0.05) were used for over-representation pathway analysis using the clusterProfiler R package and WikiPathways database.

Data integration: Integration was performed on the donor and recipient datasets independently using the following workflow in Seurat. Normalization via SCTransform was performed for each time point separately. The SelectIntegrationFeatures function was then used to find consistently variable features across datasets. We identified anchors which were used to integrate the datasets with subsequent principal component analysis, clustering, and visualization being performed on the integrated counts. For all differential gene expression testing in the integrated dataset, log normalized RNA counts were used as per the Seurat recommendations.

Trajectory analysis: Palantir in Python v3 was used to perform trajectory analysis in the recipient population. SCTransformed counts with mitochondrial counts and cell cycle state regressed out were used as the input. Monocytes were specified as the starting population and default parameters were used to infer pseudotime and entropy values.

MyD88 post-transplant day 3 reference mapping: The MyD88 post-transplant day 3 WT and KO datasets were processed using the same framework outlined in “pre-processing”. Differential Gene expression was used to annotate cell types a priori without reference mapping. SCTransform normalized data was also mapped onto the integrated recipient object using FindTransferAnchors() by projecting the PCA structure of the recipient onto the query dataset. Finally, MapQuery() was used to project the query dataset onto the recipient UMAP embedding to annotate query cell types from the recipient annotations. Reference mapped cell-type prediction scores were used to assess the robustness of label transfer.

### Statistical Analysis

Data were analyzed by using the statistical software (Prism, V9.0; GraphPad, La Jolla, CA). Kaplan-Meier survival analysis was performed with Log-rank (Mantel-Cox) with a p-value <0.05 considered statistically significant. Differences between groups were compared by using non-parametric Mann-Whitney U test. Multiple means were compared by using a one- or two-way analysis of variance with the Dunn’s test for multiple comparisons. P< 0.05 (two-sided) was indicative of a statistically significant difference. Bonferroni correction was performed when multiple hypotheses were tested. Data are presented as dot plots or box whisker plots generated in Prism. Power calculations were performed to ensure adequate sample size (n = number of animals). The exact sample size used to calculate statistical significance is stated in the appropriate figure legend.

### Data Availability

All sequencing data is freely available upon request.

### Code Availability

All scripts used for single-cell data analysis are available from GitHub (https://github.com/jamrute/2021_ACR_Kopecky_Lavine).

## Results

### Donor Macrophages Initially Persist after Heart Transplantation

To assess the dynamics of donor macrophages after heart transplantation, we transplanted B6 CX3CR1^GFP/+^ CCR2^RFP/+^ donor hearts into BALB/c recipients that received low-dose CTLA4-Ig. Donor CCR2^-^ macrophages (GFP^+^RFP^-^CD68^+^), donor CCR2^+^ macrophages (GFP^+^RFP^+^CD68^+^), and recipient macrophages (GFP^-^RFP^-^CD68^+^) were quantified in naïve hearts and 1, 3, 7, and 14 days after transplantation. There were more donor CCR2^-^ than CCR2^+^ macrophages in the heart at all timepoints (**Figure 1A-C**). Donor CCR2^-^ and CCR2^+^ macrophages persisted at early time points (days 1, 3) following transplantation. By post-transplant day 7, donor macrophages (CCR2^-^: 85 macrophages/mm^2^; CCR2^+^: 9.7 macrophages/mm^2^) were outnumbered by infiltrating recipient macrophages (704 recipient macrophages/mm^2^) (**Figure 1D**). Few donor macrophages (CCR2^-^: 1.6 macrophages/mm^2^; CCR2^+^: 1.1 macrophages/mm^2^) remained in heart grafts on post-transplant day 14.

**Figure 1:**
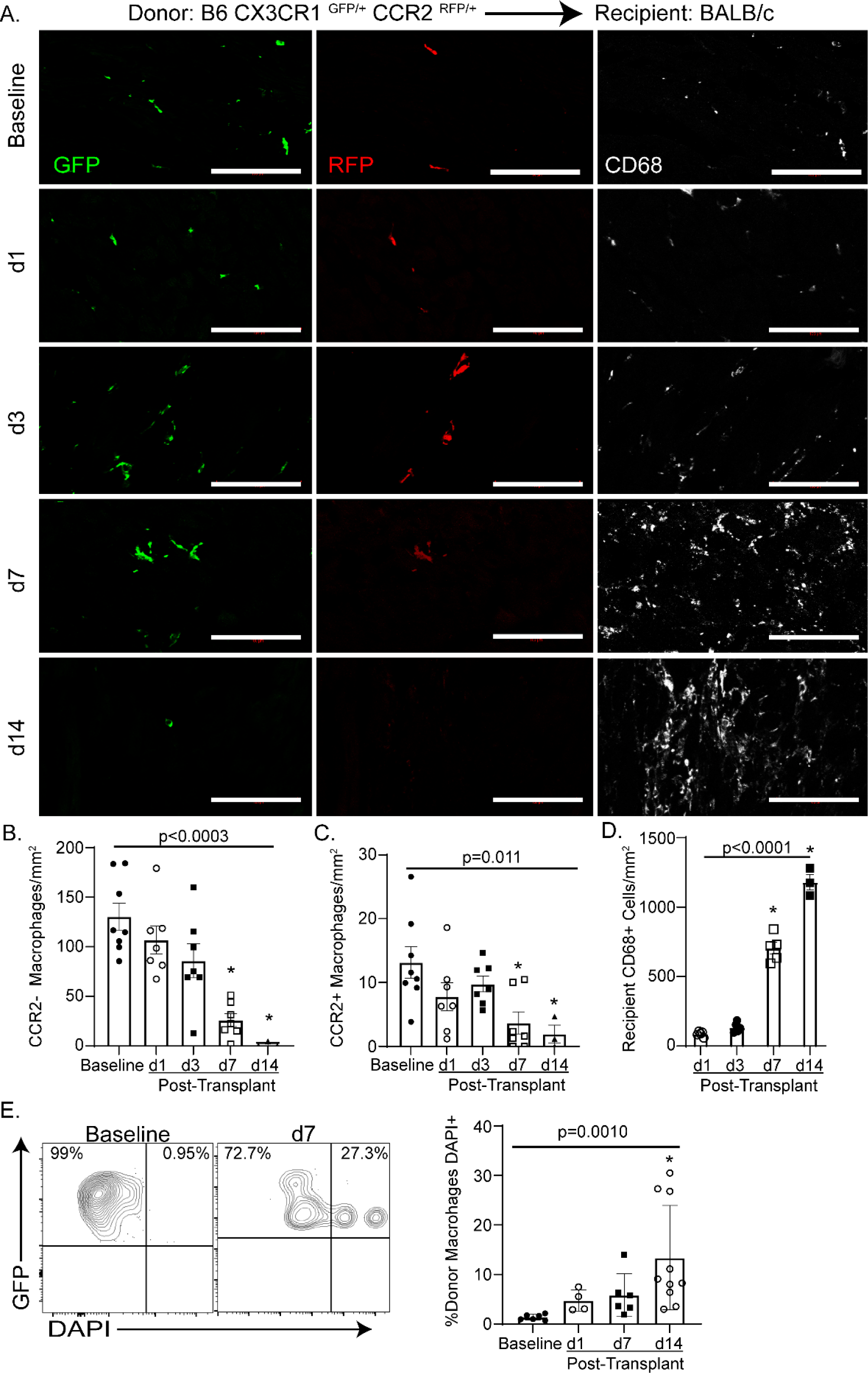
Persistence of Donor Macrophages after Heart Transplantation. A-D) At baseline, all macrophages co-express GFP and CD68. A subset of GFP^+^ CD68^+^ macrophages express RFP and are identified as donor CCR2^+^ macrophages. Donor CCR2^-^ macrophages are identified as CD68^+^ GFP^+^ RFP^-^. At post-transplant day 1 (d1), donor CCR2^-^ and CCR2^+^ macrophages co-exist with recipient derived macrophages (CD68^+^ only). By post-transplant day 14 (d14), only rare donor macrophages are identified. B) Donor CCR2^-^ macrophages significantly decrease over time from baseline to d14 (Kruskal-Wallis; p < 0.0003). Specifically, there is a decrease in donor CCR2^-^ macrophages between baseline and post-transplant day 7 (d7) (Dunn’s test for multiple comparisons; p = 0.0015) and baseline and d14 (Dunn’s test for multiple comparisons; p = 0.0014). C) There is a significant decrease of donor CCR2^+^ macrophages over time from baseline to d14 (Kruskal-Wallis; p = 0.0111) and specifically between baseline and d7 (Dunn’s test for multiple comparisons; p = 0.0197) and baseline and d14 (Dunn’s test for multiple comparisons; p = 0.0215). D) There is a significant increase in recipient CD68^+^ cells during the course of rejection (Kruskal-Wallis; p < 0.0001) and specifically between post-transplant day 1(d1) and d7 (Dunn’s test for multiple comparisons; p = 0.0134) and baseline and d14 (Dunn’s test for multiple comparisons; p = 0.0015). E) DAPI^+^ donor macrophages (GFP^+^) were considered to have undergone cell death. There was a significant increase in donor macrophages cell death after transplant (Kruskal-Wallis; p = 0.0010) and from baseline to d7 (Dunn’s test for multiple comparisons; p = 0.002) Scale bar = 100 µm.

To ensure that CCR2 expression remained stable in donor macrophages after transplantation, we performed genetic lineage tracing of CCR2^+^ cells. Tamoxifen chow was provided to B6 CCR2^ertCre/+^Rosa26^tdTomato^CCR2^GFP/+^ CD45.2 mice for two weeks prior to transplantation into CD45.1 BALB/c recipients to permanently label donor CCR2^+^ macrophages (tdTomato^+^). Flow cytometry performed 7 days after transplantation revealed that donor macrophages (CD45.2^+^CD64^+^) were either tdTomato^-^GFP^-^ (89.2%) or tdTomato^+^GFP^+^ (9.2%). Few donor macrophages were either tdTomato^-^GFP^+^ or tdTomato^+^GFP^-^ indicating that CCR2 expression is stable in donor macrophages after transplantation (**Online** **Figure 1A**).

To explore underlying mechanisms driving donor macrophage loss following transplantation, we measured donor macrophage cell death, proliferation, and emigration from the heart. Flow cytometry and TUNEL staining revealed a significant increase in donor macrophage cell death (1.3% DAPI^+^ GFP^+^ at baseline versus 13.3% DAPI ^+^ GFP ^+^ at post-transplant day14) after transplantation (p = 0.0010) (**Figure 1E, Online** **Figure 1B-E**). We next assessed both hematogenous and lymphatic emigration from the heart. Evaluation of donor macrophages (CX3CR1^GFP/+^ CCR2^RFP/+^ into BALB/c with CTLA4-Ig) within the recipient spleen revealed less than 20 donor GFP^+^ macrophages per spleen, suggestive of minimal hematogenous trafficking after transplantation (**Online** **Figure 2A**) [38]. Lymphatic connections to the heart are severed after surgery and re-establishment of lymphatic drainage from the graft may take several weeks; however, several reports identify donor immune cells trafficking to the recipient mediastinal lymph nodes [39–41]. We observed rare donor CCR2^-^ but no CCR2^+^ macrophages within the mediastinal lymph nodes (**Online** **Figure 2B**). Proliferation has been reported to contribute to cardiac macrophage persistence [22, 42]. Immunofluorescence revealed that 4-5% of donor CCR2^-^ and CCR2^+^ macrophages were proliferating (Ki67^+^) at baseline. Following transplantation, we observed an increase in the relative percentage of Ki67^+^ donor CCR2^-^ macrophages (4.3% versus 11.0% at day 7, p = 0.035) **(Online** **Figure 2C-D**). A high percentage of recipient CD68^+^ cells (5.8%) were Ki67^+^ at post-transplant day 7 (**Online** **Figure 2D**).

**Figure 2:**
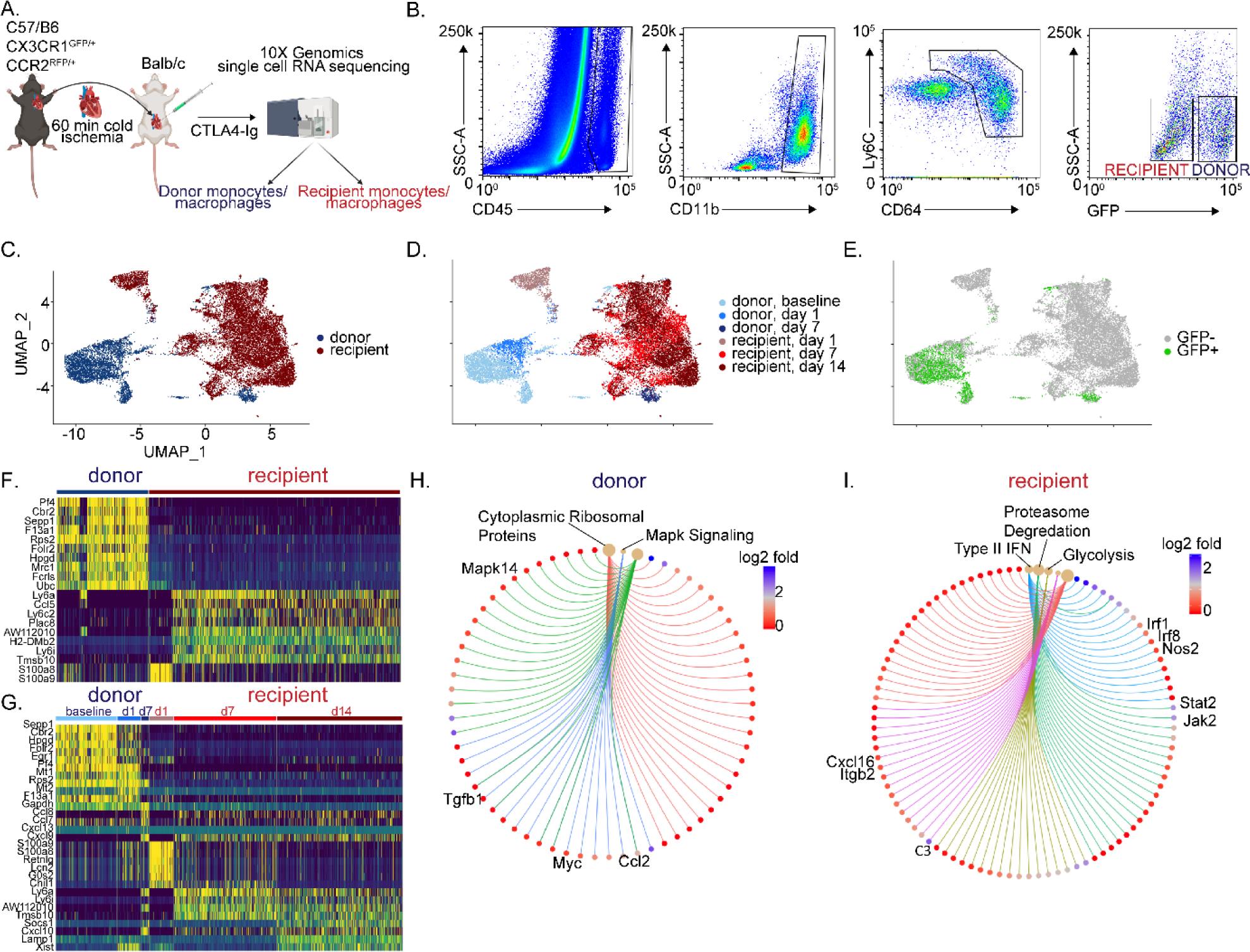
Donor and recipient immune cells are distinct. A) Schematic of single cell RNA sequencing workflow. B) Post-transplant day 1 gating scheme. Donor myeloid cells are CD45^+^ CD11b^+^ Ly6c^+^ CD64^+^ GFP^+^ whereas recipient myeloid cells are CD45^+^ CD11b^+^ Ly6c^+^ CD64^+^ GFP^-^. UMAP embedding plot of aggregated dataset split by C) donor and recipient, D) donor and recipient separated by timepoint, and E) GFP expression showing donor vs recipient demarcation, confirming robustness of gating strategy. Heatmaps of normalized counts for differentially expressed marker genes between F) donor versus recipient and G) donor versus recipient separated by timepoint. Gene-Concept Network Plot for upregulated pathways with key genes annotated in H) donors and I) recipients using statistically significant genes (adjusted p-value < 0.05) from differential gene testing.

To determine whether allograft rejection was responsible for donor macrophage cell death, we transplanted B6 CX3CR1^GFP/+^ CCR2^RFP/+^ donor hearts into BALB/c recipients and administered high-dose CTLA4-Ig [27, 43] **(Online** **Figure 2E**). This regimen has been shown to prevent cellular rejection with continued treatment but does not induce tolerance as the heart rejects upon cessation of therapy [27]. High-dose CTLA4-Ig prevented the loss of donor CCR2^-^ macrophages at post-transplant day 14. Cessation of high-dose CTLA4-Ig at post-transplant day 14 resulted in the loss of donor CCR2^-^ macrophages by day 28 post-transplant suggesting that allograft rejection leads to the elimination of donor CCR2^-^ macrophages. High-dose CTLA4-Ig did not prevent the loss of donor CCR2^+^ macrophages (**Online** **Figure 2F**). These data indicate that rejection may contribute to the loss of CCR2^-^, but not CCR2^+^ donor macrophages.

### Donor and Recipient Macrophages are Distinct and Evolve Over Time after Heart Transplantation

We utilized single cell RNA sequencing to investigate the cellular and transcriptional landscape of donor and recipient macrophages after heart transplantation. B6 CX3CR1^GFP/+^ CCR2^RFP/+^ donor hearts were transplanted into BALB/c recipients and received low-dose CTLA4-Ig (**Figure 2A**). From the transplanted heart, donor (CD45^+^CD11b^+^CD64^+^GFP^+^) and recipient (CD45^+^CD11b^+^CD64^+^<u>GFP</u>^-^) cells were sorted at baseline (donor) and 1 (donor and recipient), 7 (donor and recipient) and 14 (recipient) days after transplantation and underwent single cell RNA sequencing (**Figure 2B**). We recovered approximately 30,000 donor and 30,000 recipient cells across all time points and detected 3000-5000 genes per cell (**Online** **Figure 3**). Donor and recipient cells clustered separately and were readily distinguished based on GFP expression (**Figure 2C-E**). Differential gene expression analysis identified discrete transcriptional signatures within donor and recipient populations. Donor cells expressed markers of tissue resident macrophages including Cbr2, F13a1, Folr2, and Mrc1. Recipient cells expressed Ly6c2, Ly6a, Plac8, Ccl5, S100a8, and S100a9 consistent with monocyte and monocyte-derived macrophage identity (**Figure 2F**). Donor and recipient macrophages showed distinct time evolving transcriptional signatures at multiple timepoints after transplantation (**Figure 2G, Online** **Figure 4A-F**). Pathways enriched in donor macrophages include MAPK, Myc signaling and ribosomal protein synthesis. In contrast, glycolysis, electron transport chain, type II interferon signaling, proteasome degradation, and TYROBP/DAP-12 signaling pathways were enriched in recipient monocytes and macrophages (**Figure 2H-I, Online** **Figure 4G-L**).

**Figure 3:**
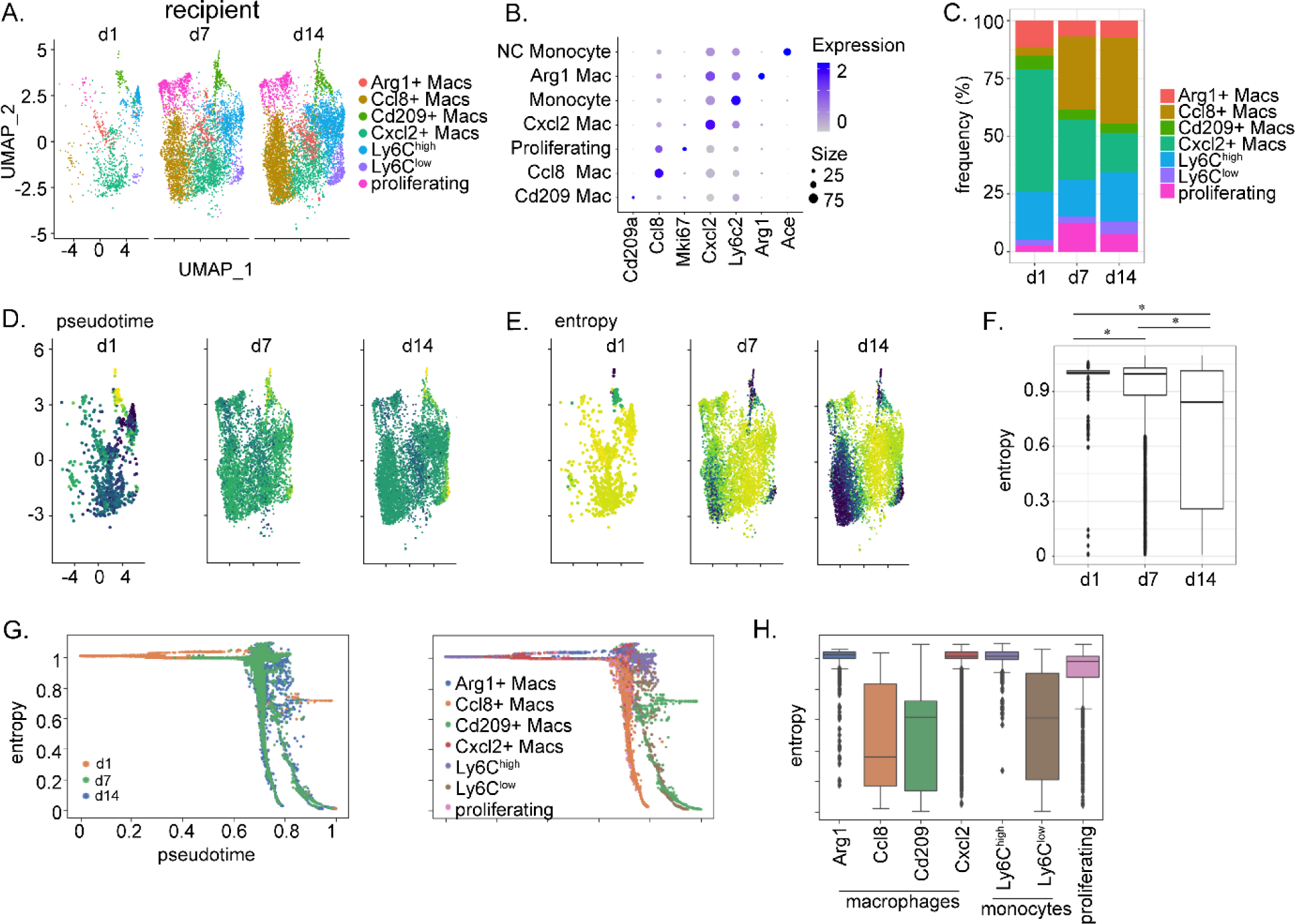
Recipient immune cells are heterogeneous and differentiate after infiltration. A) UMAP embedding plot of recipient myeloid cells at post-transplant day 1 (d1), post-transplant day 7 (d7), and post-transplant day 14 (d14) colored by cell type. B) Dot plot for top marker gene for each cell cluster found from differential gene expression testing where dot size corresponds to percentage of cells expressing the gene and color corresponds to average expression in the given cluster. C) Frequency of each population at d1, d7, and d14. Palantir D) pseudotime and E) entropy trajectories of recipient cells at d1, d7, and d14. F) Bar graph of entropy across time of aggregated clusters with p-values for associated comparisons. G) Palantir entropy versus pseudotime colored by timepoint (left) and cell type (right). H) Box plot for Palantir entropy values split by cell type.

**Figure 4:**
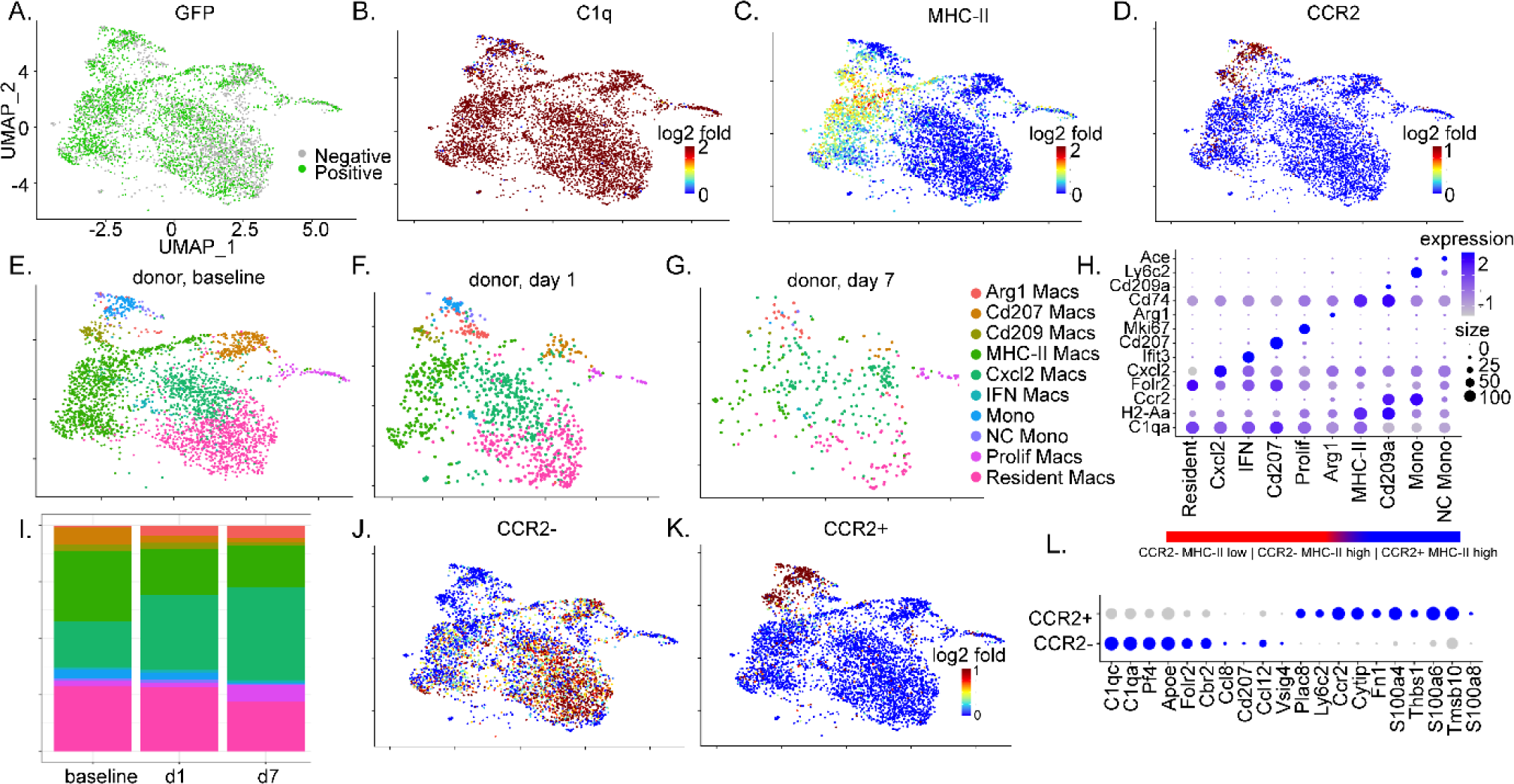
Donor macrophages can be distinguished as CCR2^+^ and CCR2^-^. UMAP embedding plot of A) GFP, B) C1qa, C) H2-Aa, and D) Ccr2 expression. UMAP embedding plot of donor macrophages colored by cluster annotation at E) baseline, F) post-transplant day 1, and G) post-transplant day 7. H) Dot plot for C1qa, H2-Aa, and Ccr2 expression and top marker genes per cell cluster found from differential gene expression testing where dot size corresponds to percentage of cells expressing the gene and color corresponds to average expression in the given cluster. I). Bar plot of relative cell composition of donor macrophages at baseline, post-transplant day 1 (d1), and post-transplant day 7 (d7). J) Feature plot of gene set scores from CCR2^-^ macrophages (C1qc, C1qa, Pf4, Apoe, Folr2, Cbr2, Ccl8, Cd207, Ccl12, Vsig4) and K) CCR2^+^ macrophages (Plac8, Ly6c2, Ccr2, Cytip, Fn1, S100a4, Thb1, S100a6, Tmsb10, S100a8). L) Dot plot for CCR2^+^ and CCR2^-^ macrophages marker genes where dot size corresponds to percentage of cells expressing the gene and color corresponds to average expression in the given cluster.

### Recipient Monocyte and Macrophage Diversity

Following transplantation, recipient monocytes infiltrate the donor heart and are known to promote allograft rejection [44]. To investigate the cell fates of infiltrating monocytes following transplantation, we performed an integrated analysis of recipient monocytes and macrophages. Cell clustering revealed distinct cell states of monocytes (Ly6C^low^, Ly6C^high^), macrophages (Arg1^+^, Cxcl2^+^, Ccl8^+^, proliferating), and dendritic-like (Cd209^+^) cells with unique transcriptional signatures (**Figure 3A-B, Online** **Figure 5**). Monocytes were prevalent across all time points after transplantation suggestive of their ongoing recruitment. Arg1^+^ and Cxcl2^+^ macrophages decreased and Ccl8^+^ macrophages increased in frequency over time suggesting a temporal relationship amongst macrophage subsets (**Figure 3C**). Palantir trajectory analysis indicated that the extent of cell entropy reduced over time after transplantation indicating that recipient monocytes and macrophages enter the heart and differentiate over the course of rejection (**Figure 3D-F**). Ly6C^high^ monocytes, Arg1^+^ macrophages, and Cxcl2^+^ macrophages had the greatest degree of entropy and lowest pseudotime values suggesting that they represent relatively undifferentiated states. In contrast, Cd209^+^ dendritic-like cells, Ly6C^low^ monocytes, and Ccl8^+^ macrophages displayed reduced entropy and higher pseudotime values suggesting that these subsets are more differentiated. Proliferating cells had intermediate entropy and pseudotime values consistent with the hypothesis that infiltrating monocytes and newly formed monocyte-derived macrophages proliferate within cardiac tissue [42] (**Figure 3G**). Analysis of the cellular differentiation landscape (entropy versus. pseudotime) predicted distinct pathways of monocyte differentiation that included Ccl8^+^ macrophage, Cd209^+^ dendritic-like cell, and Ly6C^low^ monocyte branches. Interestingly, Arg1 and Cxcl2^+^ macrophages were predicted to represent intermediate states that give rise to Ccl8^+^ macrophages (**Figure 3H, Online** **Figure 6**).

**Figure 5:**
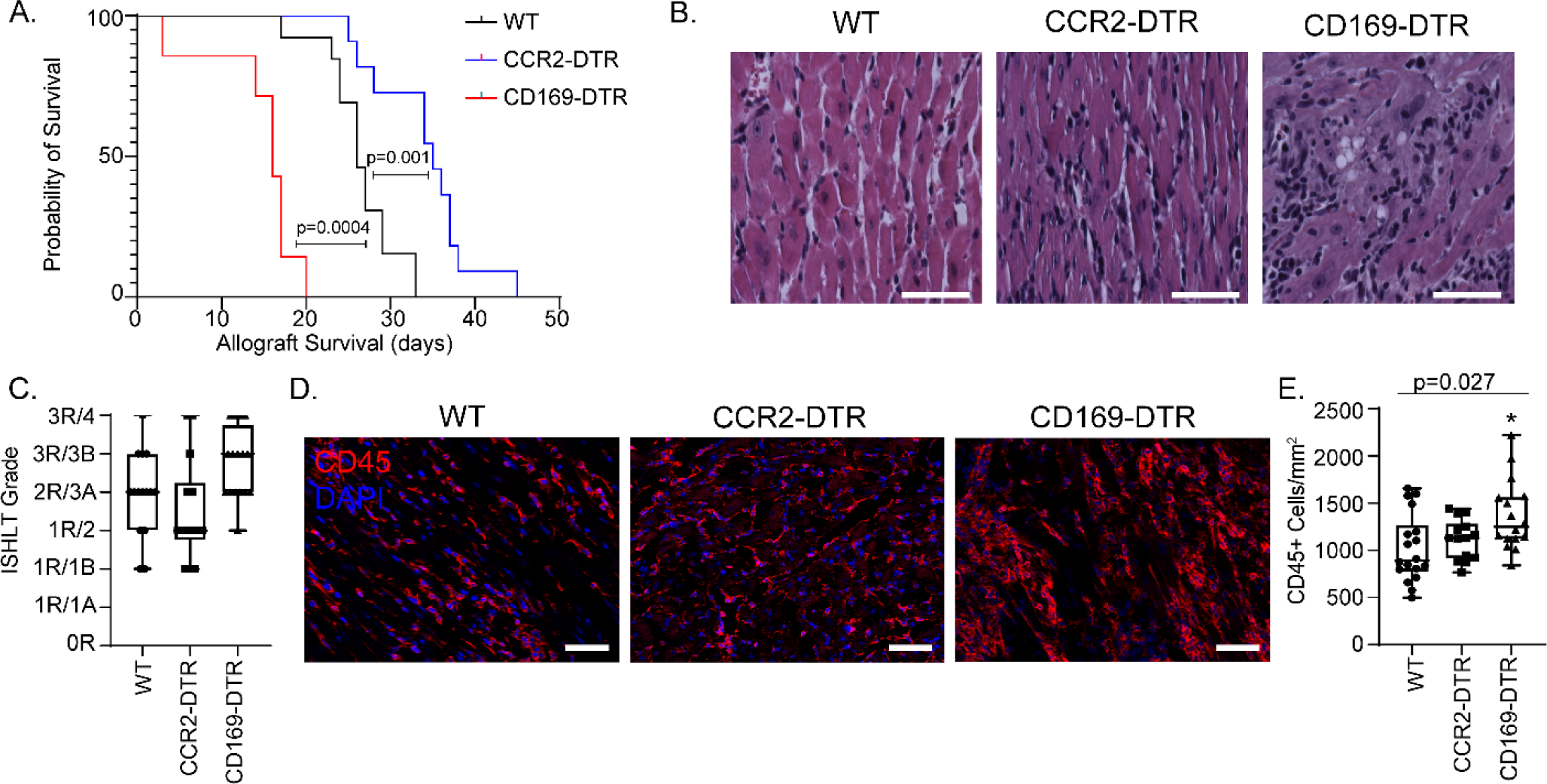
Donor macrophages differentially mediate allograft survival. A) Kaplan-Meier survival curve of CD169^DTR/+^ vs CCR2^DTR/+^ vs littermate controls (Log-rank). B) H & E stain on transplanted hearts collected at post-transplant day 10. C) At least 3 random regions were evaluated by a trained cardiac pathologist and scored based on the 1990/2004 ISHLT cellular rejection grading guidelines (n=16 hearts for each cohort). D) Post-transplant day 10 hearts were stained with CD45 (red) and DAPI (blue). E) Number of CD45^+^ cell/mm^2^ heart was quantified. There was a significant difference among the three groups (Kruskal-Wallis; p = 0.027) and between WT and CD169^DTR/+^ (Dunn’s test for multiple comparisons; p=0.014). There was no difference noted between WT and CCR2^DTR/+^ (Dunn’s test for multiple comparisons; p = 0.58). n=16 for each cohort. Scale bar = 50 µm.

**Figure 6:**
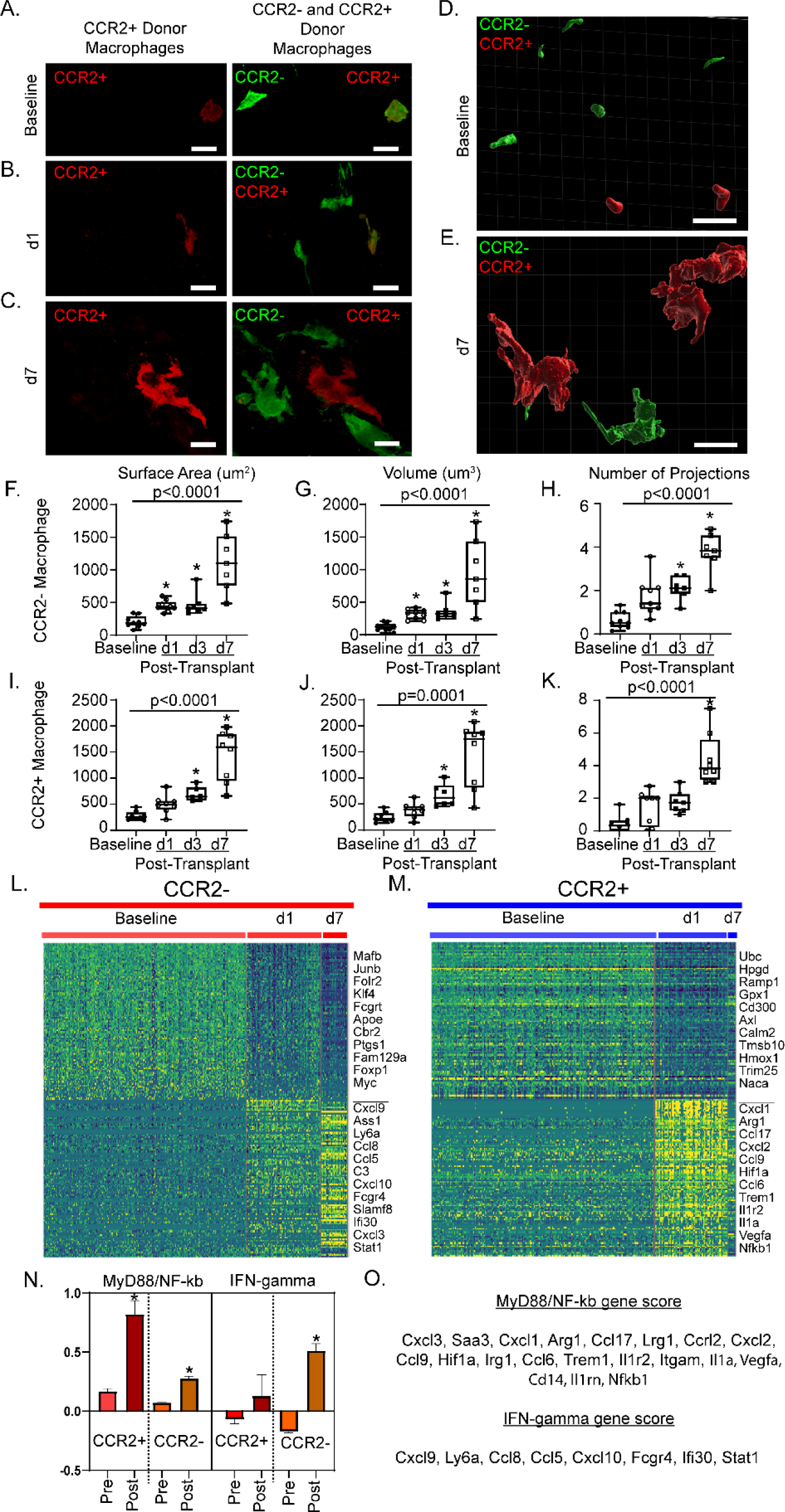
Donor macrophages dynamically respond after heart transplantation. 40X magnification z-stacks were obtained of dual reporter mice at A) baseline, B) post-transplant day 1 (d1), and C) post-transplant day 7 (d7). 30 µm thick sections were reconstructed with Imaris software to obtain volumetric reconstructions of donor macrophages at D) baseline and E) at d7. From reconstructions, quantitative measurements of donor CCR2^-^ macrophages F) surface area (µm^2^), G) volume (µm^3^) and H) number of projections were obtained. For each timepoint, each data point represents the average of 10-20 macrophages from at least two regions of interest in at least two separate sections per heart. Identical measurements were performed in donor CCR2^+^ macrophages (I-K) across time. Statistical analyses were computed with Kruskal-Wallis (noted p-value) and Dunn’s test for multiple comparisons (* when < 0.05 compared to baseline). Heatmaps of normalized counts for differentially expressed marker genes between baseline and post-transplant for L) donor CCR2^-^ macrophages and M) donor CCR2^+^ macrophages. N) Gene set scores in donor CCR2^-^ and CCR2^+^ macrophages for NF-κβ /MYD88 (CCR2^+^: p < 0.0001; CCR2^-^: p < 0.0001) and IFN-γ (CCR2 : p = 0.61; CCR2 : p < 0.0001) split by pre- and post-transplant status. Gene set scores calculated with Mann-Whitney U test. * < 0.05. O) Genes used to calculate gene set score. Scale bar = 10 µm.

### Transcriptional Diversity of Donor CCR2^-^ and CCR2^+^ Macrophages

The composition of donor macrophage populations was assayed using our single cell RNA sequencing dataset. Donor-derived macrophages were identified by GFP expression (**Figure 4A**) and expressed the macrophage marker, C1q (**Figure 4B**). We had previously categorized cardiac macrophages based on MHC-II and CCR2 expression [19, 21, 22]. Consistent with these findings, we detected MHC-II^high^ CCR2^-^, MHC-II^low^ CCR2^-^, MHC-II^high^ CCR2^+^, and MHC-II^low^ CCR2^+^ populations (**Figure 4C-D**). High-resolution clustering revealed further transcriptional diversity amongst MHC-II^low^ CCR2^-^ macrophages. We identified two transcriptional states that displayed a resident macrophage phenotype (Folr2^+^, Folr2^+^Cd207^+^Vsig4^+^) and three additional states distinguished by the expression of type I interferon responsive genes, chemokines (Cxcl2, Ccl7), and Arg1. MHC-II^high^ CCR2^+^ macrophages expressed Cd209a and MHC-II^low^ CCR2^+^ cells expressed Ly6c2 (classical monocytes) and Ace (non-classical monocytes) (**Figure 4E-H**, **Online** **Figure 7A-K**). Temporal analysis comparing donor macrophages at baseline to days 1 and 7 post-transplant revealed loss of donor macrophages over time, consistent with our immunostaining and flow cytometry data. Most donor CCR2^+^ macrophages were lost by day 7 post-transplant (**Online** **Figure 7L**). While many donor CCR2^-^ macrophage states decreased over time, MHC-II^low^ CCR2^-^ macrophages expressing Cxcl2/Ccl7 and Arg1 increased in relative abundance (**Figure 4I**).

**Figure 7:**
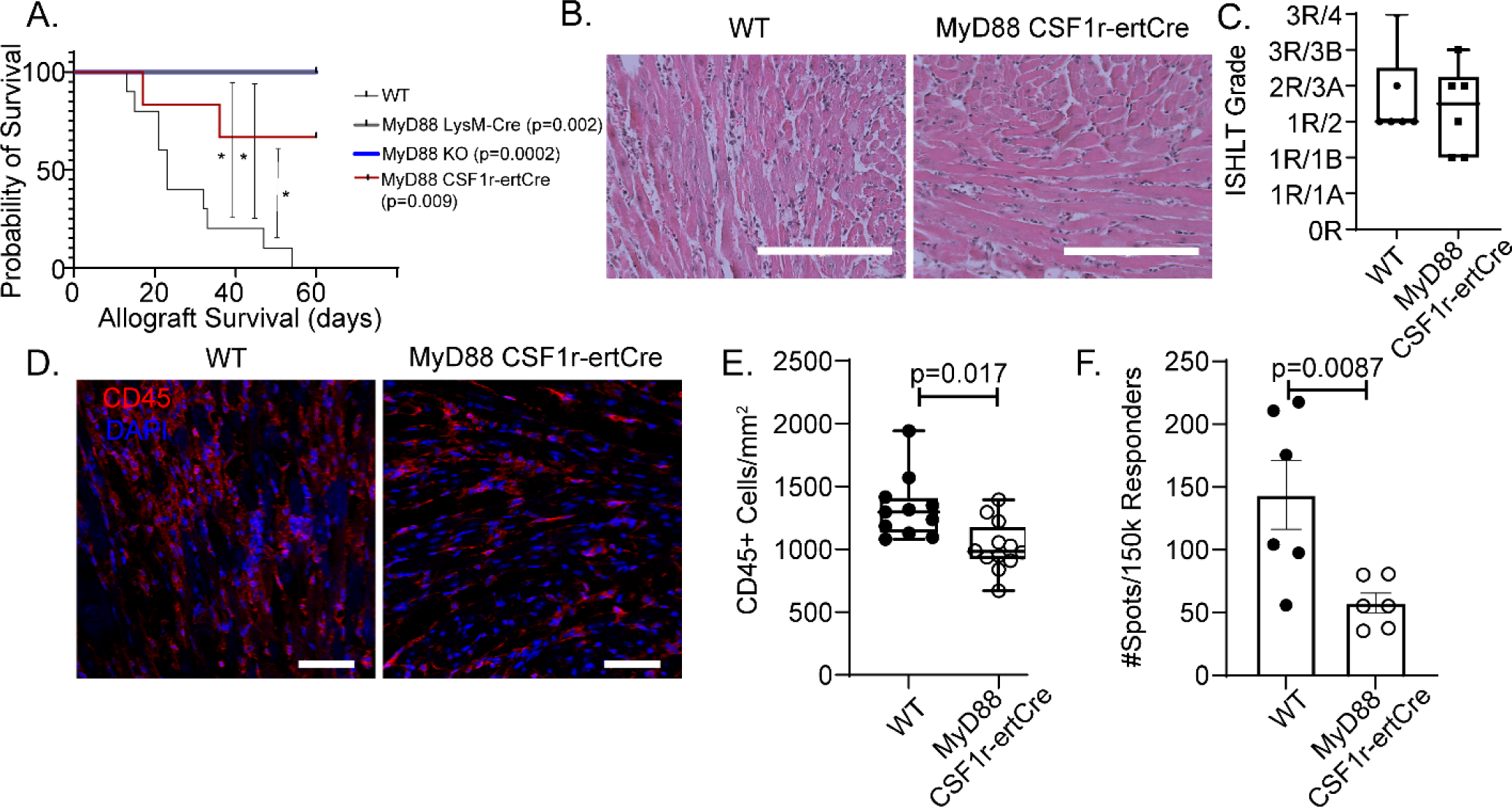
Donor macrophages signal through MyD88. A) Kaplan-Meier survival curve with control versus MyD88^f/f^ LysM^Cre/+^ (Log-rank; p = 0.002), MyD88^f/f^ CSF1r^ertCre/+^ (Log-rank; p = 0.009), or MyD88^-/-^ (“KO” Log-rank; p = 0.0002) donor allografts (censored at 60 days). B) H & E staining of WT and MyD88^f/f^ CSF1r^ertCre/+^ allografts collected at post-transplant day 10. C) ISHLT cellular rejection scores comparing WT (n=12) and MyD88^f/f^ CSF1r^ertCre/+^ (n=12) hearts. D) CD45^+^ immunofluorescent staining of WT (n=12) and MyD88^f/f^ CSF1r^ertCre/+^ (n=12) hearts with E) quantification showing significant reduction in CD45^+^ cells/mm^2^ in MyD88^f/f^ CSF1r^ertCre/+^ hearts (Mann-Whitney U Test; p = 0.0017,) compared to littermate control. F) Number of IFN-γ spots per 150,000 responder T-cells with a significant reduction in MyD88^f/f^ CSF1r^ertCre/+^ cohort (Mann-Whitney U Test; p = 0.0087). Scale bar: B= 200 µm; D=50 µm.

To delineate transcriptomic differences between donor CCR2^-^ and CCR2^+^ macrophages, we performed a differential expression analysis. Donor CCR2^-^ macrophages differentially expressed tissue resident macrophages genes including Folr2, Cbr2, Vsig4, Ccl8, and Ccl12. In contrast, donor CCR2^+^ macrophages expressed inflammatory genes (**Figure 4J-L**). Collectively, these data indicate that donor CCR2^-^ and donor CCR2^+^ macrophages represent distinct populations at the transcriptional level.

### Donor Macrophages are Essential for Allograft Survival after Heart Transplantation

To elucidate the functional roles of donor macrophages in allogenic heart transplantation, we utilized CD169^DTR/+^ [28] and CCR2^DTR/+^ [29] mice to deplete CCR2^-^ and CCR2^+^ macrophages from the cardiac graft, respectively. We have previously established the specificity and efficacy of these murine strains in depleting CCR2^-^ and CCR2^+^ macrophages from the heart [19, 24]. To this end, we transplanted B6 control, CD169^DTR/+^, or CCR2^DTR/+^ hearts into BALB/c recipients that received low-dose CTLA4-Ig. Diphtheria toxin (DT) (200 ng) was administered to the donor mouse daily for three days prior to transplant and three times per week to recipient after transplantation to maintain depletion. Allografts lacking donor CCR2^+^ macrophages (CCR2^DTR/+^) had longer survival than littermate controls (35 versus. 26 days, p=0.0013; log-rank HR 0.287, 0.079-1.031). In contrast, allografts lacking donor CCR2^-^ macrophages (CD169^DTR/+^) had shorter survival times compared to littermate controls (16 versus. 28 days, p=0.0004; log-rank HR 4.334, 1.147-16.39) (**Figure 5A**).

To investigate whether donor macrophages influence allograft inflammation, we harvested donor hearts 10 days after transplantation and performed histological analysis (**Figure 5B-C**). Compared to controls, allografts lacking donor CCR2^-^ macrophages had more severe cellular rejection (3R/3B versus 2R/3A), while allografts lacking donor CCR2^+^ macrophages had milder cellular rejection (1R/2 versus 2R/3A). Immunostaining revealed increased CD45^+^ leukocytes in allografts lacking donor CCR2^-^ macrophages compared to controls (1367 versus 1021 CD45^+^ cells/mm^2^, p = 0.014). Allografts lacking donor CCR2^+^ macrophages had a similar number of CD45^+^ leukocytes compared to controls (1116 versus 1021 CD45^+^ cells/mm^2^, p = 0.58) (**Figure 5D-E**). These data support the conclusion that donor CCR2^-^ macrophages confer protection against acute allograft rejection whereas donor CCR2^+^ macrophages promote allograft rejection.

### Donor Macrophage Activation following Transplantation

To examine the phenotypic behavior of donor macrophages after heart transplantation, we performed high-resolution imaging of donor macrophages at baseline and 1, 3, and 7 days post-transplant **(Figure 6A-C, Online** **Figure 8A-B**). At baseline, donor CCR2^-^ and CCR2^+^ macrophages were relatively small, homogenously distributed within the heart, and displayed few cellular projections. Three-dimensional reconstruction of 30 µm heart sections (**Figure 6D-E**), revealed that both donor CCR2^-^ and CCR2^+^ macrophages increase significantly in surface area (CCR2^-^: baseline 202.6 µm^2^ versus day 7 1121 µm^2^, p<0.0001; CCR2^+^: baseline 277.6µm^2^ versus day 7 1431 µm^2^, p<0.0001), volume (CCR2^-^: baseline 120.1 µm^3^ versus day 7 941.9 µm^3^, p<0.0001; CCR2^+^: baseline 245.1 µm^3^ versus day 7 1431 µm^3^, p<0.0001, and number of projections (CCR2^-^: baseline 0.637 projections/macrophage versus day 7 3.76 projections/macrophage, p<0.0001; CCR2^+^: baseline 0.43 projections/macrophage versus day 7 4.388 projections/macrophage, p<0.0001) (**Figure 6F-K**). There was no significant difference in morphologic quantification between donor macrophages at each time point (p > 0.05), except for CCR2^-^ macrophage volume at baseline compared to that of CCR2^+^ macrophages (p = 0.006) (**Online** **Figure 8C-E**).

**Figure 8:**
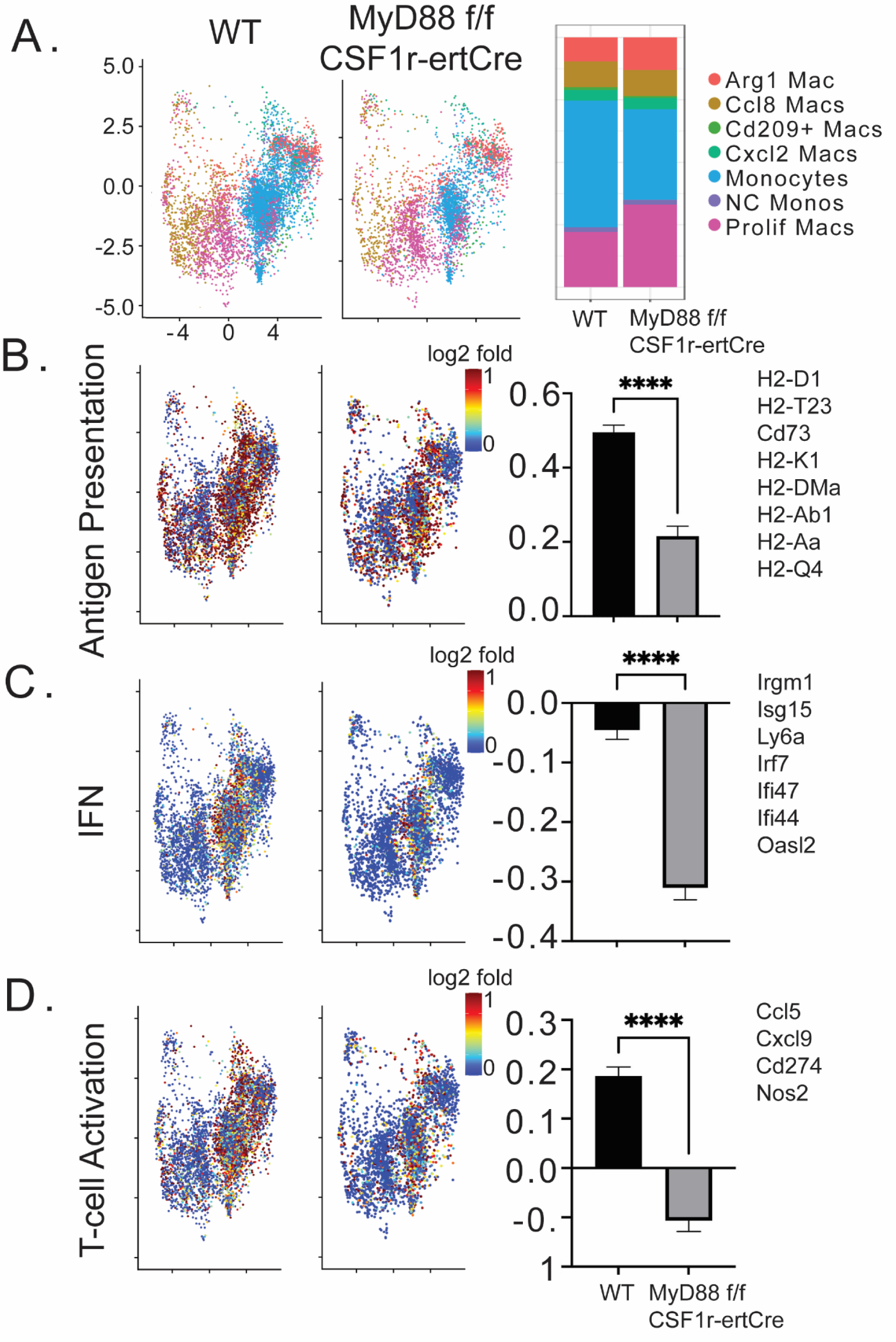
Donor MyD88 depletion leads to modulation of recipient immune cell gene expression. UMAP embedding plot of A) post-transplant day 3 WT and post-transplant day 3 MyD88^f/f^ CSF1r^ertCre/+^ mapped onto recipient composite dataset (left). Composition plot showing frequency of each population (right) in WT and MyD^f/f^ CSF1r^ertCre/+^. Combined z-scores for top genes and calculated gene set scores for B) antigen presentation (0.569 vs 0.211, p<0.0001), C) interferon signaling (-0.0032 vs -0.324, p<0.0001), and D) T-cell activation (0.1744 vs -0.1803 p< 0.0001) gene set scores with genes on the right. Statistical test Mann-Whitney U Test.

Differential gene expression analysis demonstrated that donor CCR2^-^ and CCR2^+^ macrophages displayed distinct activation signatures after transplant. Compared to baseline, donor CCR2^+^ macrophages expressed markedly increased levels of genes associated with enhanced MYD88/NF-signaling post-transplant (0.17 versus 0.82, p < 0.0001) (**Figure 6M, O**). Donor CCR2^-^ macrophages also displayed a subtle MYD88/NF-κβ signature after transplantation (0.072 versus 0.28, p < 0.0001). Donor CCR2^-^ macrophages differentially expressed genes involved in type II interferon signaling, specifically interferon gamma (IFN-γ) (-0.17 versus 0.51, p < 0.0001) (**Figure 6L, N-O**). Taken together, these data indicate that both donor CCR2^-^ and CCR2^+^ macrophages are activated through distinct mechanisms following transplantation.

### Inhibition of MYD88 Signaling in Donor Macrophages Prevents Allograft Rejection

We have previously demonstrated that donor CCR2^+^ macrophages orchestrate leukocyte trafficking following syngeneic transplantation through a MYD88 dependent mechanism [19, 24]. To determine whether inhibition of MYD88 signaling in donor macrophages presents a possible therapeutic approach for preventing or reducing graft rejection, we used donor mice that lack MyD88 either globally or selectively in macrophages. We transplanted B6 control, MyD88^-/-^, MyD88^f/f^ LysM^Cre/+^, or MyD88^f/f^ CSF1r^ertCre/+^donor hearts into BALB/c recipients which were treated with low-dose CTLA4-Ig. Global deletion of MYD88 in the donor heart and conditional deletion of MYD88 in donor macrophages both resulted in significantly prolonged allograft survival compared to controls (all p < 0.009) (**Figure 7A**).

To assess graft inflammation, we performed histological analyses and CD45^+^ immunostaining on control and MyD88^f/f^ CSF1r^ertCre/+^donor hearts 10 days after transplantation. Compared to controls, allografts lacking MYD88 in donor macrophages had only mild cellular rejection (1R/2 versus 2R/3A) (**Figure 7B-C**) and decreased CD45^+^ cells abundance (1022 versus 1328 CD45^+^ cells/mm^2^, p = 0.017) (**Figure 7D-E**). Furthermore, compared to controls, recipients transplanted with MyD88^f/f^ CSF1r^ertCre/+^ hearts had significantly fewer T-cells in their spleens that produced IFN-γ following exposure to donor antigen (57.4 versus 143.6 spots/150,000 responder T-cells, p = 0.0087). (**Figure 7F**). Collectively, these findings indicate that MYD88 signaling in donor macrophages is an important driver of alloimmunity after transplantation.

### Deletion of MYD88 effects Antigen Presentation, Interferon Signaling, and T-cell Activation

Depletion of MyD88 resulted in prolonged allograft survival, reduced inflammation, and reduced allograft alloreactivity. Our single cell RNA sequencing data suggests that infiltrating monocytes differentiate into distinct cell types after transplantation. To delineate the role of donor macrophages influenced recipient monocyte fate specification, we utilized a MyD88 donor macrophage depletion strategy. We performed single cell RNA sequencing of recipient monocytes, macrophages, and dendritic cells (CD45^+^ CD11b^+^ Ly6c^+/-^ CD64^+/-^) isolated from WT and MyD88^f/f^ CSF1r^ertCre/+^ donor hearts at post-transplant day 3. We recovered approximately 20,000 cells and detected 3,000 to 5,000 genes per cell (**Online Figure 9A**). The majority of recovered cells expressed markers of monocytes and macrophages present within the recipient dataset including Arg1^+^ and Ccl8^+^ cluster marker genes (**Online Figure 9B-D**). We also observed a population of MHC-II^high^ macrophages that were reduced in MyD88^f/f^ CSF1r^ertCre/+^ donor hearts. Few Lyve1^+^ resident macrophages were identified consistent with a predominance of recipient derived cells.

To examine the impact of donor macrophage MYD88 signaling on recipient cell fate, we projected the data onto our recipient monocyte, macrophage, and dendritic cell reference object and obtained high confidence mapping scores (**Figure 8A, Online Figure 9E**). Cell composition analysis showed increased proportion of monocytes in WT allografts and increased proliferating cells in MyD88^f/f^ CSF1r^ertCre/+^ donor hearts. We did not detect any differences among other monocyte, macrophage, or dendritic cell states (**Figure 8A**). However, we did detect global differences in gene expression. MyD88^f/f^ CSF1r^ertCre/+^ cells displayed lower expression of genes associated with antigen presentation (0.22 versus 0.50, p < 0.001) (**Figure 8B**), interferon signaling (-3.11 versus -0.045, p < 0.001) (**Figure 8C**), and T-cell activation (-0.107 versus 0.187, p < 0.001) (**Figure 8D**). These data are consistent with the observation that deletion of MyD88 in donor macrophages reduces alloreactive T-cell generation and suggestive of decreased antigen presentation capacity in infiltrating recipient macrophages as a potential mechanism.

## Discussion

To the best of our knowledge, this is the first study to investigate the transcriptional landscape and dynamics of donor and recipient myeloid cells following allogenic heart transplantation at a single cell resolution. We defined the functional importance of donor CCR2^+^ and CCR2^-^ macrophages in allograft rejection and identified donor CCR2^+^ macrophages as a key cell type that potentiates rejection. Finally, we implicated MYD88 signaling in donor macrophages as a potential therapeutic target to reduce the generation of alloreactive T-cells, decrease rejection, and extend allograft survival.

Using genetic lineage tracing and single cell RNA sequencing, we longitudinally dissected the dynamics of donor macrophages after heart transplantation. We show that in the setting of cellular rejection, donor macrophages persisted for approximately 1 week and are subsequently lost from the graft. Intriguingly, suppression of allograft rejection using high-dose CTLA4-Ig immunosuppression was sufficient to preserve donor CCR2^-^ macrophages, but not donor CCR2^+^ macrophages. This observation raises the possibility that alloreactive T-cells play an active role in eliminating donor CCR2^-^ macrophages. Future studies will be required to identify the precise population of T-cells that target donor CCR2^-^ macrophages and to dissect the molecular mechanisms and cell death pathways involved.

Single cell RNA sequencing demonstrated that donor and recipient macrophages are transcriptionally distinct and that these populations dynamically evolve over the course of rejection. Consistent with transcriptional evidence of cell activation, donor macrophages show significant morphologic changes including increased surface area, volume, and cellular projections. Differential gene expression analysis identified signatures of IFN-γ and MYD88/NF-κβ signaling in donor CCR2^-^ and CCR2^+^ macrophages, respectively, after transplantation. Either depletion of donor CCR2^+^ macrophages or conditional deletion of MYD88 signaling in donor macrophages was sufficient to reduce rejection and extend allograft survival. These data build on our prior observation that donor CCR2^+^ macrophages direct infiltration of recipient neutrophils and monocytes into the heart through the expression of MYD88 dependent chemokines and cytokines [24]. Depletion of donor macrophage MYD88 signaling led to infiltration of recipient cells with lower expression of antigen presentation, interferon, and T-cell activation gene expression signatures. Based on the robust MYD88/NF-κβ signature observed in donor CCR2^+^ macrophages after transplantation and improved outcomes seen in donor hearts that lack MyD88 in macrophages, we posit that MYD88 signaling in donor CCR2^+^ macrophages contributes to allograft rejection. However, we cannot exclude the possibility that MYD88 signaling in donor CCR2^-^ macrophages may also contribute to some of the observed phenotypes. Collectively, these findings provide the first evidence that targeting specific donor macrophage populations may serve as a therapeutic approach to prevent cellular rejection. Targeting donor specific populations could be done using *ex-situ* perfusion devices prior to implantation and holds promise of ameliorating graft rejection [45, 46].

Our data indicated an opposing role for donor CCR2^-^ macrophages. Depletion of donor CCR2^-^ macrophages resulted in rapid rejection and increased immune cell infiltration into the allograft, indicating that donor CCR2^-^ macrophages have a protective function. Following transplantation, we observed patches of donor CCR2^-^ macrophages that appeared to physically interact with and clustered around donor CCR2^+^ macrophages. It is possible that donor CCR2^-^ macrophages prevent the activation of donor CCR2^+^ macrophages and other immune cells. Consistent with this hypothesis, tissue resident macrophages have previously been shown to cloak areas of injury and prevent inflammatory activation of innate immune cells [47]. The mechanistic basis for this phenomenon is not fully understood, but could involve IFN-γ signaling. Along these lines, we hypothesize that donor CCR2 macrophage are protective, potentially through IFN-γ, and their depletion could lead to more rapid rejection and increased inflammatory infiltration. IFN-γ could be involved in the upregulation of immunomodulatory mediators.

Finally, our single cell RNA sequencing data revealed that donor and recipient macrophages are distinct and shed new light on the surprising diversity of recipient monocyte, macrophage, and dendritic cell populations. Using a probabilistic model to predict the trajectories of differentiating monocytes, we show evidence that recipient monocytes have the capacity to differentiate into several unique macrophage and dendritic cell-like populations. The environmental cues and molecular signals responsible for these fate decisions are of considerable interest. Future studies will focus on the contribution of each recipient monocyte, macrophage, and dendritic cell population to allograft rejection.

### Limitations and Conclusions

Our study is not without limitations. Mouse models of heterotopic heart transplantation are imperfect as they are mechanically unloaded and thus may not fully mirror the clinical scenario. We additionally recognize genetic manipulation of donor macrophage composition and signaling may not be fully recapitulated by pharmacological interventions. Nonetheless, our findings establish donor macrophages as a potential therapeutic target to improve transplant outcomes and identify inhibition of MYD88 signaling in the donor heart as a therapeutic approach to suppress donor CCR2^+^ macrophage activation and blunt alloimmune responses.

## Acknowledgements

We thank Geetika Bajpai and Inessa Lokshina for technical expertise and mouse husbandry.

## Sources of Funding

B.K. was supported by the Principles in Cardiovascular Research Training Grant (T32 HL007081) and the Washington University Physician Scientist Training Program. K.L. is supported by funding provided from the NHLBI (R01 HL138466, R01HL139714, R01 HL151078), Leducq Foundation Network (#20CVD02), Burroughs Welcome Fund (1014782), Children’s Discovery Institute of Washington University and St. Louis Children’s Hospital (CH-II-2015-462, CH-II-2017-628, PM-LI-2019-829), and Foundation of Barnes-Jewish Hospital (8038-88). D.K. is supported by NIH (P01AI116501 and R01 HL094601), Veterans Administration Merit Review (1I01BX002730), and the Foundation for Barnes-Jewish Hospital. We acknowledge the McDonnel Genome Institute, DDRCC histology core (P30 DK52574), and the Washington University Center for Cellular Imaging (WUCCI), which is supported in part by Washington University School of Medicine, The Children’s Discovery Institute of Washington University and St. Louis Children’s Hospital (CDI-CORE-2015-505 and CDI-CORE-2019-813) and the Foundation for Barnes-Jewish Hospital (3770) for access to imaging resources and valuable technical assistance.

## Disclosures

The authors have nothing to disclose.

## Notes

### Competing Interest Statement

The authors have declared no competing interest.

## References

1. Colvin, M., et al., OPTN/SRTR 2016 Annual Data Report: Heart. Am J Transplant, 2018. 18 Suppl 1: p. 291-362.

2. Moayedi, Y., et al., Survival Outcomes After Heart Transplantation: Does Recipient Sex Matter? Circ Heart Fail, 2019. 12(10): p. e006218.

3. Chang, D.H., M.M. Kittleson, and J.A. Kobashigawa, Immunosuppression following heart transplantation: prospects and challenges. Immunotherapy, 2014. 6(2): p. 181–94.

4. Alba, A.C., et al., Complications after Heart Transplantation: Hope for the Best, but Prepare for the Worst. International Journal of Transplantation Research and Medicine, 2016. 2(2).

5. McDonald-Hyman, C., L.A. Turka, and B.R. Blazar, Advances and challenges in immunotherapy for solid organ and hematopoietic stem cell transplantation. Sci Transl Med, 2015. 7(280): p. 280rv2.

6. Millington, T.M. and J.C. Madsen, Innate immunity and cardiac allograft rejection. Kidney Int Suppl, 2010(119): p. S18–21.

7. Kopecky, B.J., et al., Role of donor macrophages after heart and lung transplantation. Am J Transplant, 2020. 20(5): p. 1225–1235.

8. Zhang, W., N. Egashira, and S. Masuda, Recent Topics on The Mechanisms of Immunosuppressive Therapy-Related Neurotoxicities. Int J Mol Sci, 2019. 20(13).

9. Costanzo, M.R., et al., The International Society of Heart and Lung Transplantation Guidelines for the care of heart transplant recipients. J Heart Lung Transplant, 2010. 29(8): p. 914–56.

10. Halloran, P.F., Immunosuppressive drugs for kidney transplantation. N Engl J Med, 2004. 351(26): p. 2715–29.

11. Hunt, S.A. and F. Haddad, The changing face of heart transplantation. J Am Coll Cardiol, 2008. 52(8): p. 587–98.

12. Costello, J.P., T. Mohanakumar, and D.S. Nath, Mechanisms of chronic cardiac allograft rejection Tex Heart Inst J, 2013. 40(4): p. 395–9.

13. Conde, P., et al., DC-SIGN(+) Macrophages Control the Induction of Transplantation Tolerance. Immunity, 2015. 42(6): p. 1143–58.

14. Schetters, S.T.T., et al., Mouse DC-SIGN/CD209a as Target for Antigen Delivery and Adaptive Immunity. Front Immunol, 2018. 9: p. 990.

15. van den Bosch, T.P., et al., Targeting the Monocyte-Macrophage Lineage in Solid Organ Transplantation. Front Immunol, 2017. 8: p. 153.

16. Li, J., et al., The Evolving Roles of Macrophages in Organ Transplantation. J Immunol Res, 2019. 2019: p. 5763430.

17. Tse, G.H. and J. Hughes, Macrophages and transplant rejection: a novel future target? Transplantation, 2013. 96(11): p. 946–8.

18. Chen, S., F.G. Lakkis, and X.C. Li, The many shades of macrophages in regulating transplant outcome. Cell Immunol, 2020. 349: p. 104064.

19. Bajpai, G., et al., Tissue Resident CCR2- and CCR2+ Cardiac Macrophages Differentially Orchestrate Monocyte Recruitment and Fate Specification Following Myocardial Injury. Circ Res, 2019. 124(2): p. 263–278.

20. Ochando, J., et al., Trained immunity in organ transplantation. Am J Transplant, 2020. 20(1): p. 10–18.

21. Lavine, K.J., et al., Distinct macrophage lineages contribute to disparate patterns of cardiac recovery and remodeling in the neonatal and adult heart. Proc Natl Acad Sci U S A, 2014. 111(45): p. 16029–34.

22. Epelman, S., K.J. Lavine, and G.J. Randolph, Origin and functions of tissue macrophages. Immunity, 2014. 41(1): p. 21–35.

23. Bajpai, G., et al., The human heart contains distinct macrophage subsets with divergent origins and functions. Nat Med, 2018. 24(8): p. 1234–1245.

24. Li, W., et al., Heart-resident CCR2(+) macrophages promote neutrophil extravasation through TLR9/MyD88/CXCL5 signaling. JCI Insight, 2016. 1(12).

25. Pietra, B.A., et al., CD4 T cell-mediated cardiac allograft rejection requires donor but not host MHC class II. J Clin Invest, 2000. 106(8): p. 1003–10.

26. Corry, R.J., H.J. Winn, and P.S. Russell, Primarily vascularized allografts of hearts in mice. The role of H-2D, H-2K, and non-H-2 antigens in rejection. Transplantation, 1973. 16(4): p. 343-50.

27. Schwarz, C., et al., The Immunosuppressive Effect of CTLA4 Immunoglobulin Is Dependent on Regulatory T Cells at Low But Not High Doses. Am J Transplant, 2016. 16(12): p. 3404–3415.

28. Miyake, Y., et al., Critical role of macrophages in the marginal zone in the suppression of immune responses to apoptotic cell-associated antigens. J Clin Invest, 2007. 117(8): p. 2268–78.

29. Hohl, T.M., et al., Inflammatory monocytes facilitate adaptive CD4 T cell responses during respiratory fungal infection. Cell Host Microbe, 2009. 6(5): p. 470–81.

30. Hou, B., B. Reizis, and A.L. DeFranco, Toll-like receptors activate innate and adaptive immunity by using dendritic cell-intrinsic and -extrinsic mechanisms. Immunity, 2008. 29(2): p. 272–82.

31. Clausen, B.E., et al., Conditional gene targeting in macrophages and granulocytes using LysMcre mice. Transgenic Res, 1999. 8(4): p. 265–77.

32. Qian, B.Z., et al., CCL2 recruits inflammatory monocytes to facilitate breast-tumour metastasis. Nature, 2011. 475(7355): p. 222-5.

33. Adachi, O., et al., Targeted disruption of the MyD88 gene results in loss of IL-1- and IL-18-mediated function. Immunity, 1998. 9(1): p. 143–50.

34. Jung, S., et al., Analysis of fractalkine receptor CX(3)CR1 function by targeted deletion and green fluorescent protein reporter gene insertion. Mol Cell Biol, 2000. 20(11): p. 4106–14.

35. Saederup, N., et al., Selective chemokine receptor usage by central nervous system myeloid cells in CCR2-red fluorescent protein knock-in mice. PLoS One, 2010. 5(10): p. e13693.

36. Truett, G.E., et al., Preparation of PCR-quality mouse genomic DNA with hot sodium hydroxide and tris (HotSHOT). Biotechniques, 2000. 29(1): p. 52, 54.

37. Stewart, S., et al., Revision of the 1990 working formulation for the standardization of nomenclature in the diagnosis of heart rejection. J Heart Lung Transplant, 2005. 24(11): p. 1710-20.

38. Larsen, C.P., P.J. Morris, and J.M. Austyn, Donor dendritic leukocytes migrate from cardiac allografts into recipients’ spleens. Transplant Proc, 1990. 22(4): p. 1943–4.

39. Edwards, L.A., et al., Chronic Rejection of Cardiac Allografts Is Associated With Increased Lymphatic Flow and Cellular Trafficking. Circulation, 2018. 137(5): p. 488–503.

40. Soong, T.R., et al., Lymphatic injury and regeneration in cardiac allografts. Transplantation, 2010. 89(5): p. 500–8.

41. Brown, K., et al., SPECT/CT lymphoscintigraphy of heterotopic cardiac grafts reveals novel sites of lymphatic drainage and T cell priming. Am J Transplant, 2011. 11(2): p. 225–34.

42. Sager, H.B., et al., Proliferation and Recruitment Contribute to Myocardial Macrophage Expansion in Chronic Heart Failure. Circ Res, 2016. 119(7): p. 853–64.

43. Young, J.S., et al., Successful Treatment of T Cell-Mediated Acute Rejection with Delayed CTLA4-Ig in Mice. Front Immunol, 2017. 8: p. 1169.

44. Oberbarnscheidt, M.H., et al., Non-self recognition by monocytes initiates allograft rejection. J Clin Invest, 2014. 124(8): p. 3579–89.

45. Chew, H.C., P.S. Macdonald, and K.K. Dhital, The donor heart and organ perfusion technology. J Thorac Dis, 2019. 11(Suppl 6): p. S938–S945.

46. Wang, L., et al., Ex situ heart perfusion: The past, the present, and the future. J Heart Lung Transplant, 2021. 40(1): p. 69–86.

47. Uderhardt, S., et al., Resident Macrophages Cloak Tissue Microlesions to Prevent Neutrophil-Driven Inflammatory Damage. Cell, 2019. 177(3): p. 541–555 e17.

