## Supplemental Figures for "Donor Macrophages Modulate Rejection after Heart Transplantation"

SUPPLEMENTAL MATERIAL

Online Figure 1:

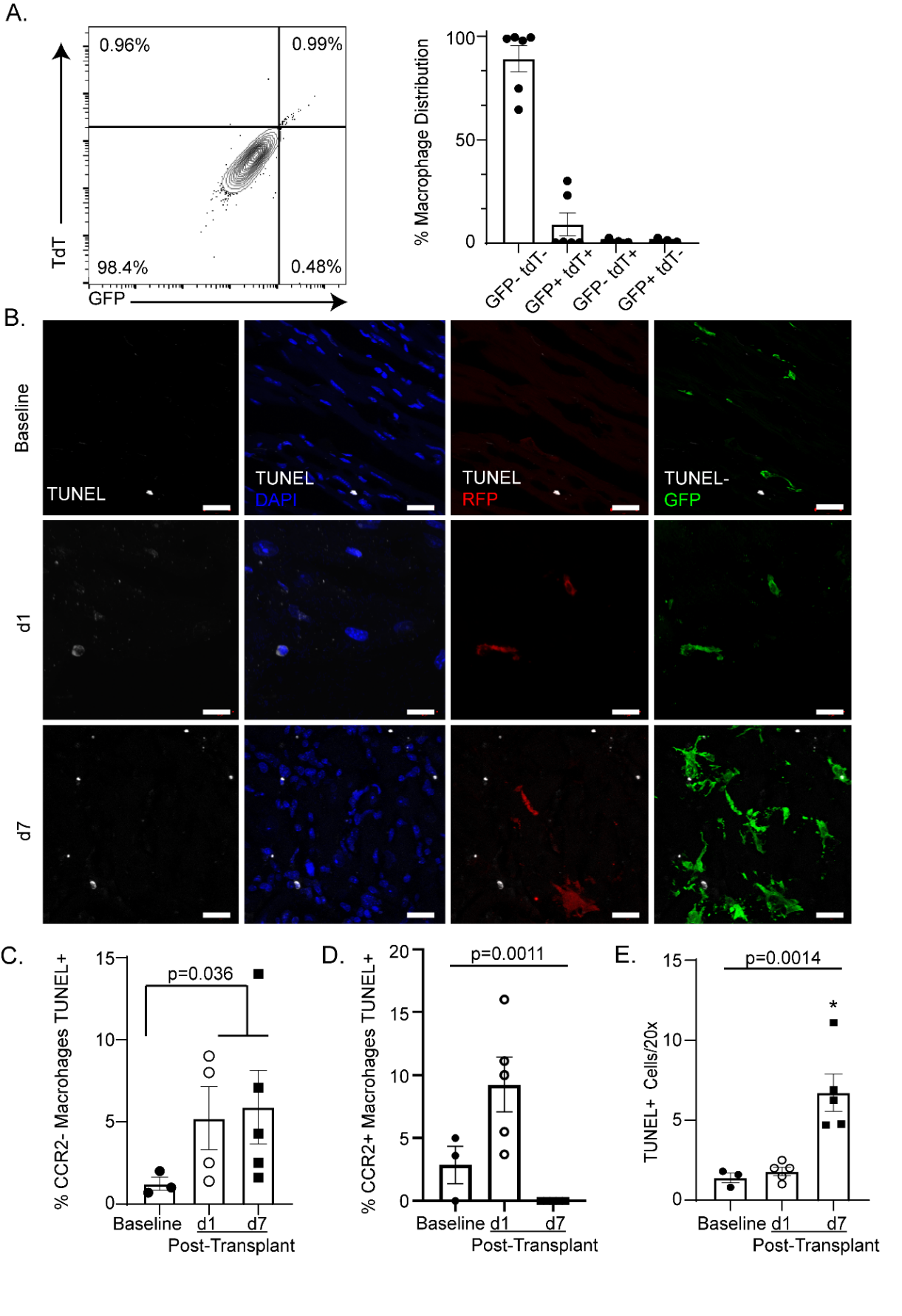

**Online Figure 1:** A) B6 CCR2^ertCre/+^ Rosa26^tdTomato^ CCR2^GFP/+^ CD45.2 mice were transplanted into CD45.1 BALB/c recipients and post-transplant day 7 hearts were gated on CD45.2^+^ CD11b^+^ Ly6c^-^ CD64^+^ and assayed for double positive (GFP^+^ tdT^+^), double negative (GFP^-^ tdT^-^), or single positive (GFP^+^ or tdT^+^). Quantification showed the majority of donor macrophages were CCR2^-^ (GFP^-^ TdT^-^; 89.17%, n=6 hearts) or CCR2^+^ (9.19%), with a minority expressing tdT^+^ but not GFP (0.75%) or expressing GFP only (0.87%) (Kruskal-Wallis; p = 0.0031). B) TUNEL immunofluorescence of donor CCR2^+^ and CCR2^-^ macrophages at baseline, post-transplant day 1 (d1), and post-transplant day 7 (d7) showing TUNEL (column 1), TUNEL and DAPI (column 2), TUNEL and RFP (column 3), and TUNEL and GFP (column 4). C) The percentage of donor CCR2^-^ macrophages that are TUNEL^+^ increase after transplant (baseline versus combined d1 and d7; Mann Whitney U Test; p = 0.0364). D) The percentage of donor CCR2^+^ macrophages that are TUNEL^+^ increase after transplant (Kruskal-Wallis; p = 0.0011). There were few donor CCR2^+^ macrophages remaining at d7. E) Quantification of total TUNEL^+^ cells (DAPI and TUNEL^+^) per 20X field at baseline, d1, and d7 show increased cell death (Kruskal-Wallis; p = 0.0014) and baseline versus d7 (Dunn’s test for multiple comparisons; p = 0.0139). Scale bar = 20 µm.

Online Figure 2:

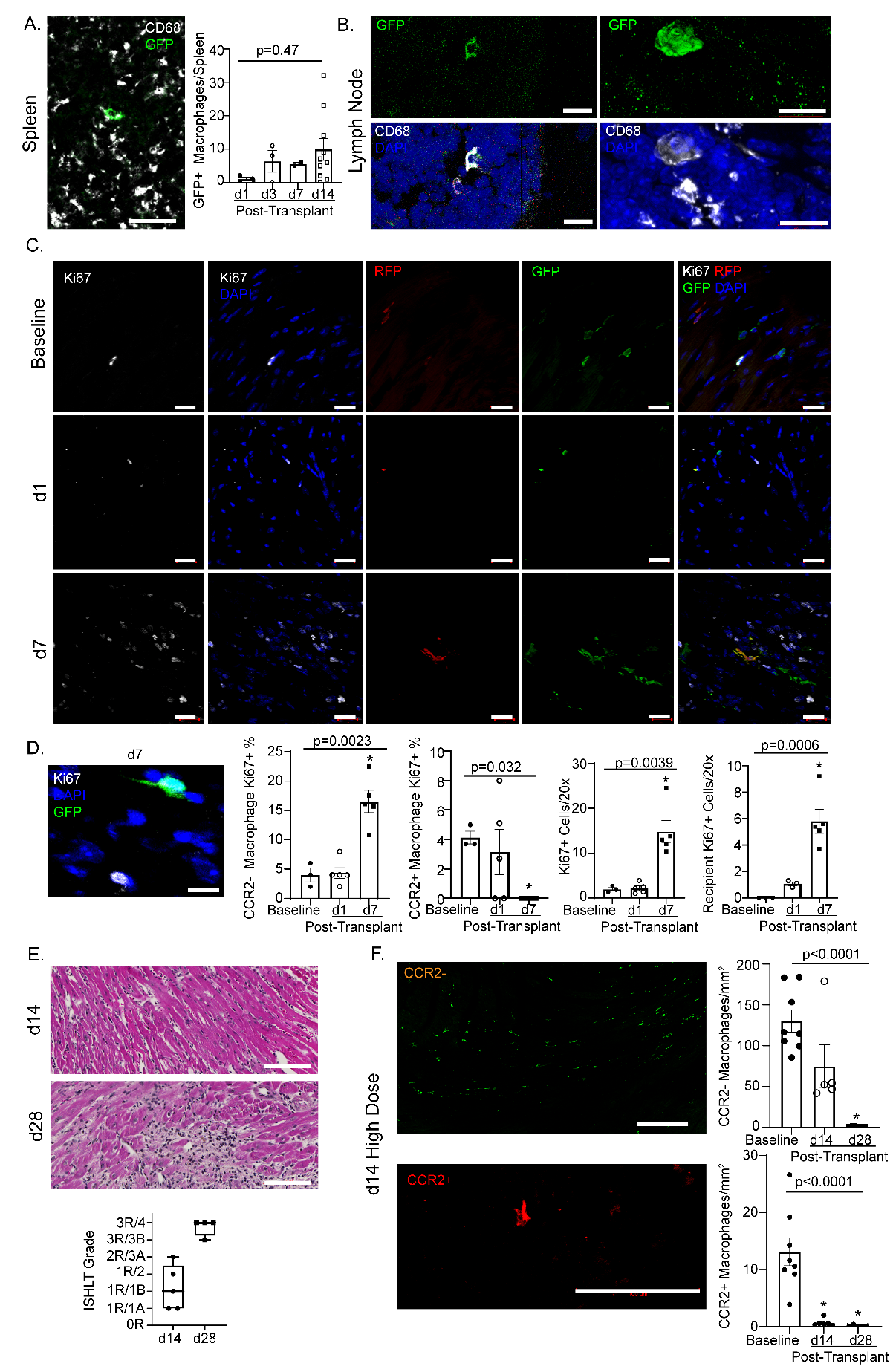

**Online Figure 2:** A) Immunofluorescent staining of donor CCR2^-^ (CD68^+^ GFP^+^) macrophage in the recipient spleen. Quantification of donor macrophages by flow cytometry which showed no significant increase in donor macrophage (GFP^+^) accumulation in the spleen after between post-transplant day 1 (d1) and post-transplant day 14 (d14) (Kruskal-Wallis; p = 0.47). B) Identification of rare donor CCR2^-^ (CD68^+^ GFP^+^) macrophage in recipient lymph node. C-D) Donor CCR2^-^ and CCR2^+^ macrophages that co-labelled with Ki67^+^ were considered proliferating. A greater percentage of CCR2^-^ macrophages were proliferating after transplant (Kruskal-Wallis; p = 0.0023) and specifically at post-transplant day 7 (d7) compared to baseline (Dunn’s test for multiple comparison; p = 0.034). The percentage of proliferating donor CCR2^+^ macrophages decreased after transplant (Kruskal-Wallis; p = 0.032) and was significantly decreased at d7 (Dunn’s test for multiple comparison; p = 0.042) compared to baseline. After transplant, there was an overall increase in proliferation of all cells (DAPI + Ki67) (Kruskal-Wallis; p = 0.0039) and significantly at d7 (Dunn’s test for multiple comparison; p = 0.0382) compared to baseline. Recipient CD68^+^ cells showed increased proliferation after transplant (Kruskal-Wallis; p = 0.0006) and specifically at d7 (Dunn’s test for multiple comparison; p = 0.0071) compared to baseline. E) High-dose CTLA4-Ig was administered until d14. At d14, heart grafts were generally scored as mild cellular rejection (1R); however, grafts collected at post-transplant day 28 (d28) were scored as severe cellular rejection (3R). F) At d14, similar numbers of donor CCR2^-^ macrophages were identified compared to baseline (Dunn’s test for multiple comparison; p = 0.26) with a significant reduction after cessation of CTLA4-Ig (d14 versus d28; Dunn’s test for multiple comparison; p = 0.0008). Compared to baseline, there was a significant loss of donor CCR2^+^ macrophages at both d14 (Dunn’s test for multiple comparison; p = 0.031) and at d28 (Dunn’s test for multiple comparison; p = 0.0003). Scale bar: A =40 µm, B-C = 10 µm, D = 20 µm, E = 100 µm, F (top) = 200 µm, F (bottom) = 100 µm.

Online Figure 3:

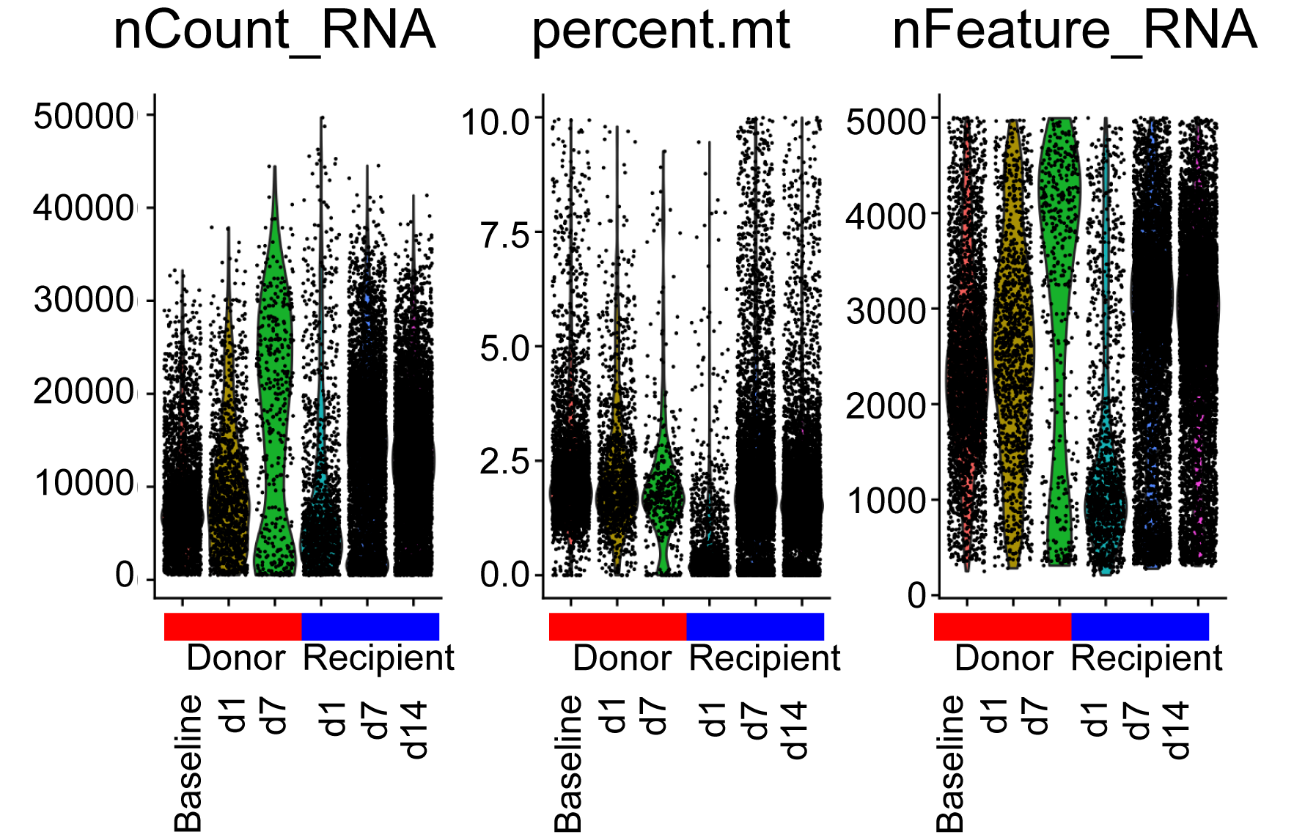

Quality control data plots showing RNA count, percent mitochondria, and feature counts of each population at baseline, post-transplant day 1 (d1), post-transplant day 7 (d7), and post-transplant day 14 (d14).

Online Figure 4:

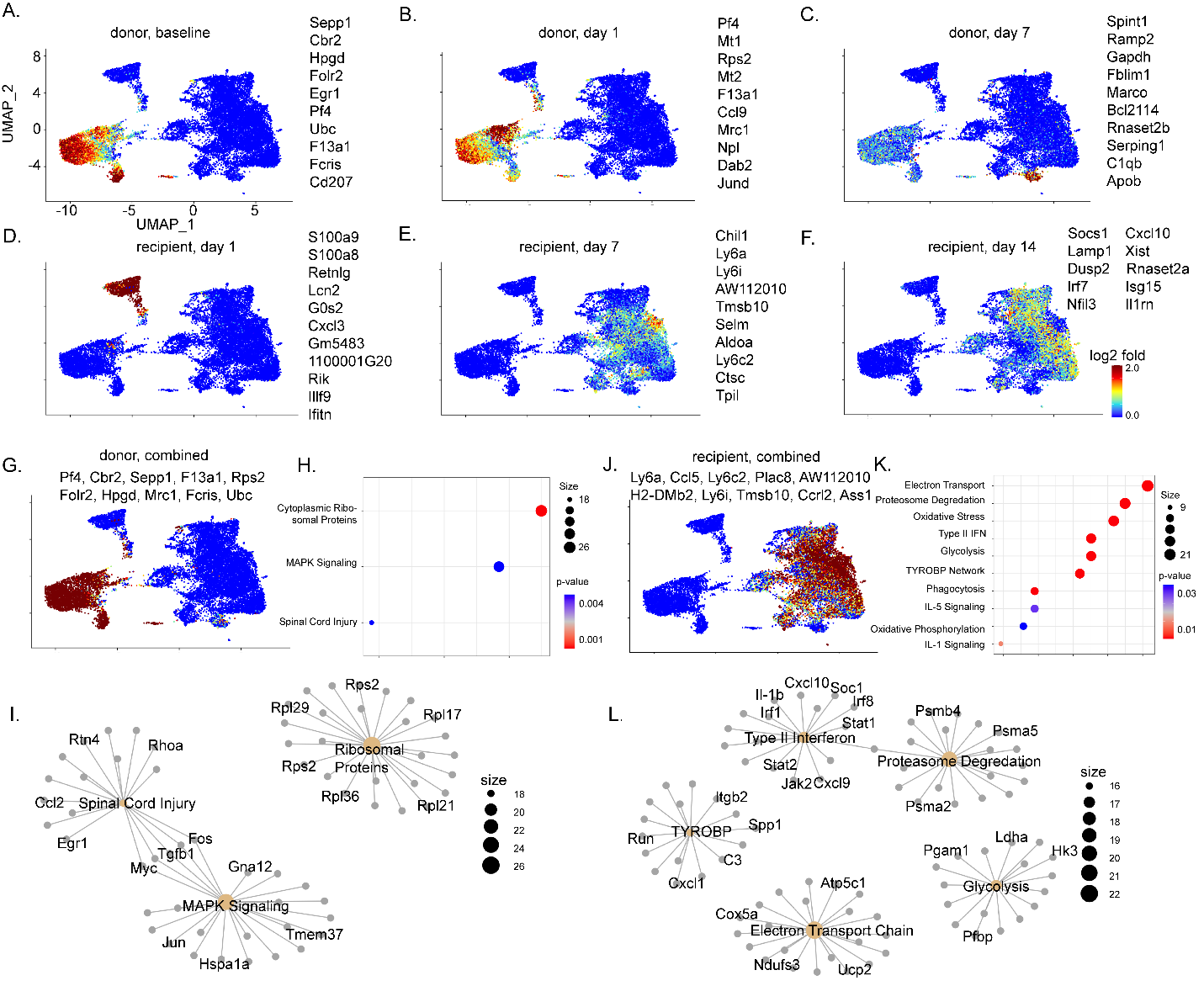

**Online Figure 4:** Combined z-score for top 10 statistically significant differentially expressed genes overlaid on UMAP embedding plot for A) donor, baseline (Sepp1, Cbr2, Hpgd, Folr2, Egr1, Pf4, Ubc, F13a1, Fcris, CD207), B) donor, post-transplant day 1 (Pf4, Mt1, Rps2, Mt2, F13a1, Ccl9, Mrc1, Npl, Dab2, Jund), C) donor, post-transplant day 7 (Spint1, Ramp2, Gapdh, Fblim1, Marco, Bcl2l14, Rnaset2b, Serping1, C1qb, Apob), D) recipient, post-transplant day 1 (S100a9, S100a8, Retnlg, Lcn2, G0s2, Cxcl3, Gm5483, 1100001G20Rik, Il1f9, Ifitn), E) recipient, post-transplant day 7 (Chil1, Ly6a, Ly6i, AW112010, Tmsb10, Selm, Aldoa, Ly6c2, Ctsc, Tpi1), F) recipient, post-transplant day 14 (Socs1, Cxcl10, Lamp1, Xist, Dusp2, Rnaset2a, Irf7, Isg15, Nfil3, Il1rn), G) combined donor (Pf4, Cbr2, Sepp1, F13a1, Rps2, Folr2, Hpgd, Mrc1, Fcris, Ubc), and J) combined recipient (Ly6a, Ccl5, Ly6c2, Plac8, AW112010, H2-DMb2, Ly6i, Tmsb10, Ccrl2, Ass1). WikiPathway over-representation analysis using statistically significant differentially expressed genes in H) donor and K) recipient. Gene-Concept Network Plot for enriched pathways with associated genes in I) donor and L) recipient.

Online Figure 5:

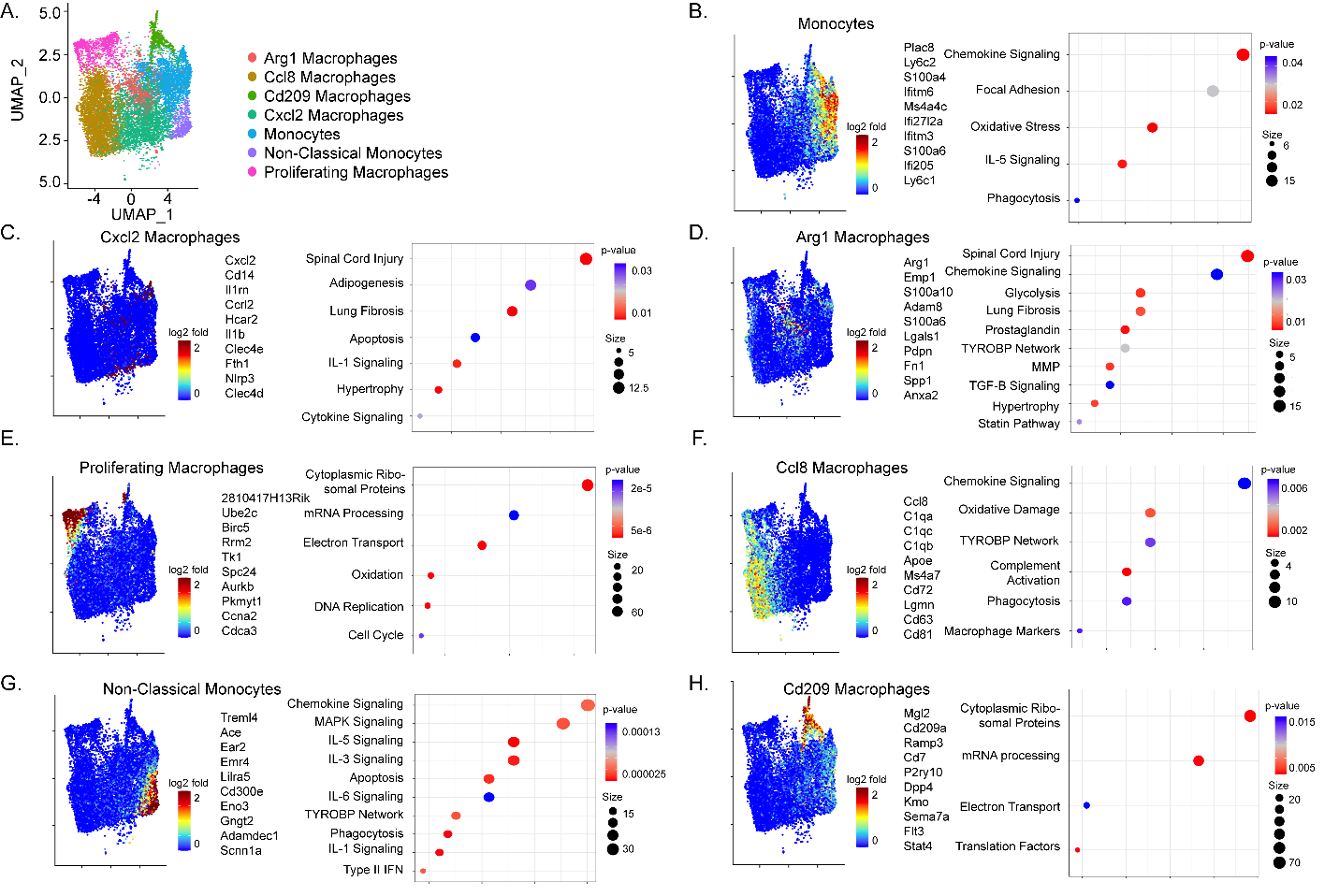

**Online Figure 5:** A) UMAP embedding plot for recipients combined across all time points colored by cell type. Combined z-score for top statistically significant differentially expressed genes overlaid on UMAP embedding plot (left) with over-representation pathway analysis using WikiPathway database (right) in B) monocytes, C) Cxcl2 macrophages, D) Arg1 macrophages, E) proliferating macrophages, F) Ccl8 macrophages, G) non-classical monocytes, and H) Cd209 macrophages.

Online Figure 6:

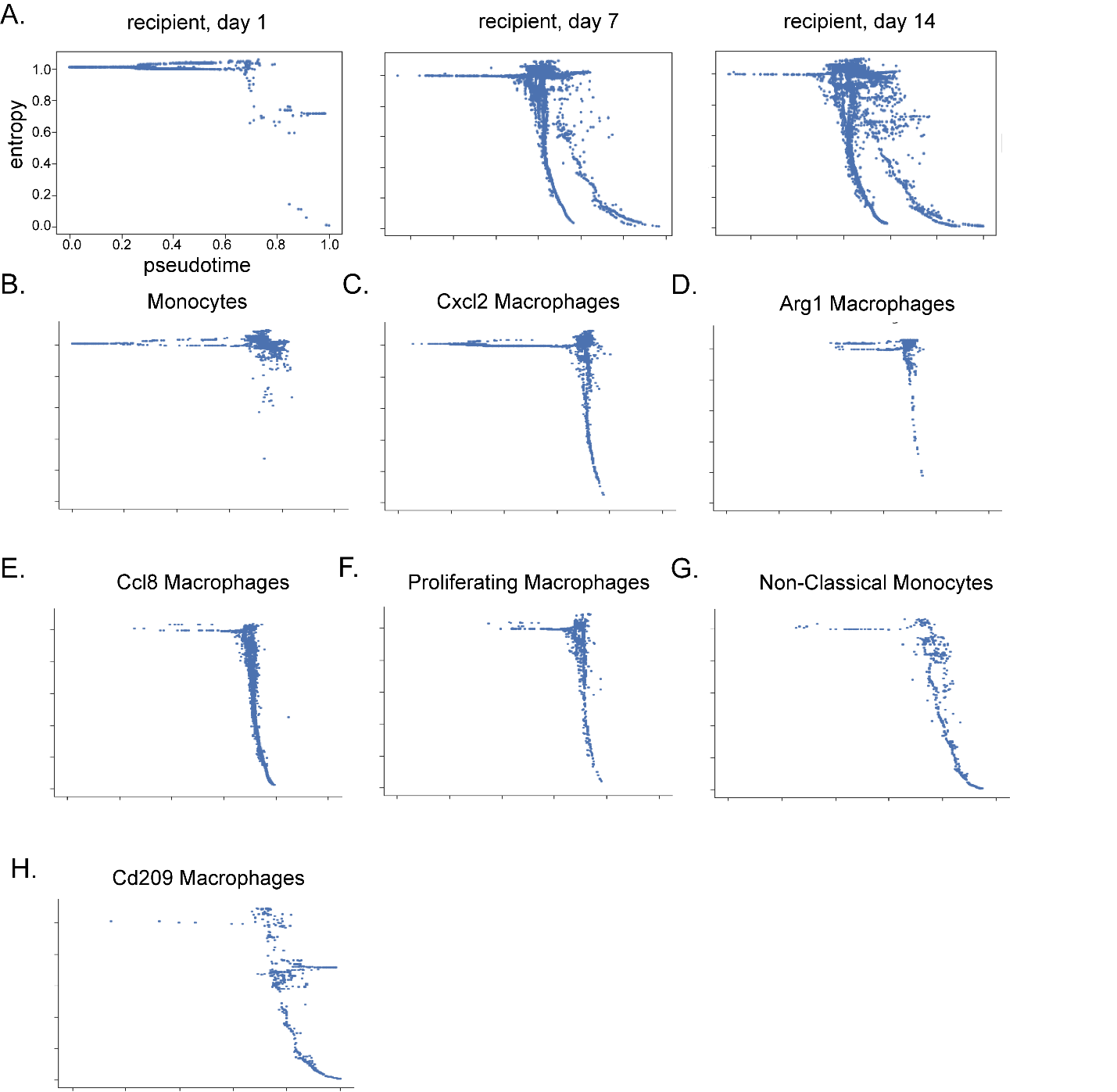

**Online Figure 6:** Palantir entropy versus pseudotime trajectories split by recipient population A) at post-transplant days 1, 7, and 14, and split by cell type B) Monocytes, C) Cxcl2 Macrophages, D) Arg1 Macrophages, E) Ccl8 Macrophages, F) Proliferating Macrophages, G) Non-classical Monocytes, and H) Cd209 Macrophages.

Online Figure 7:

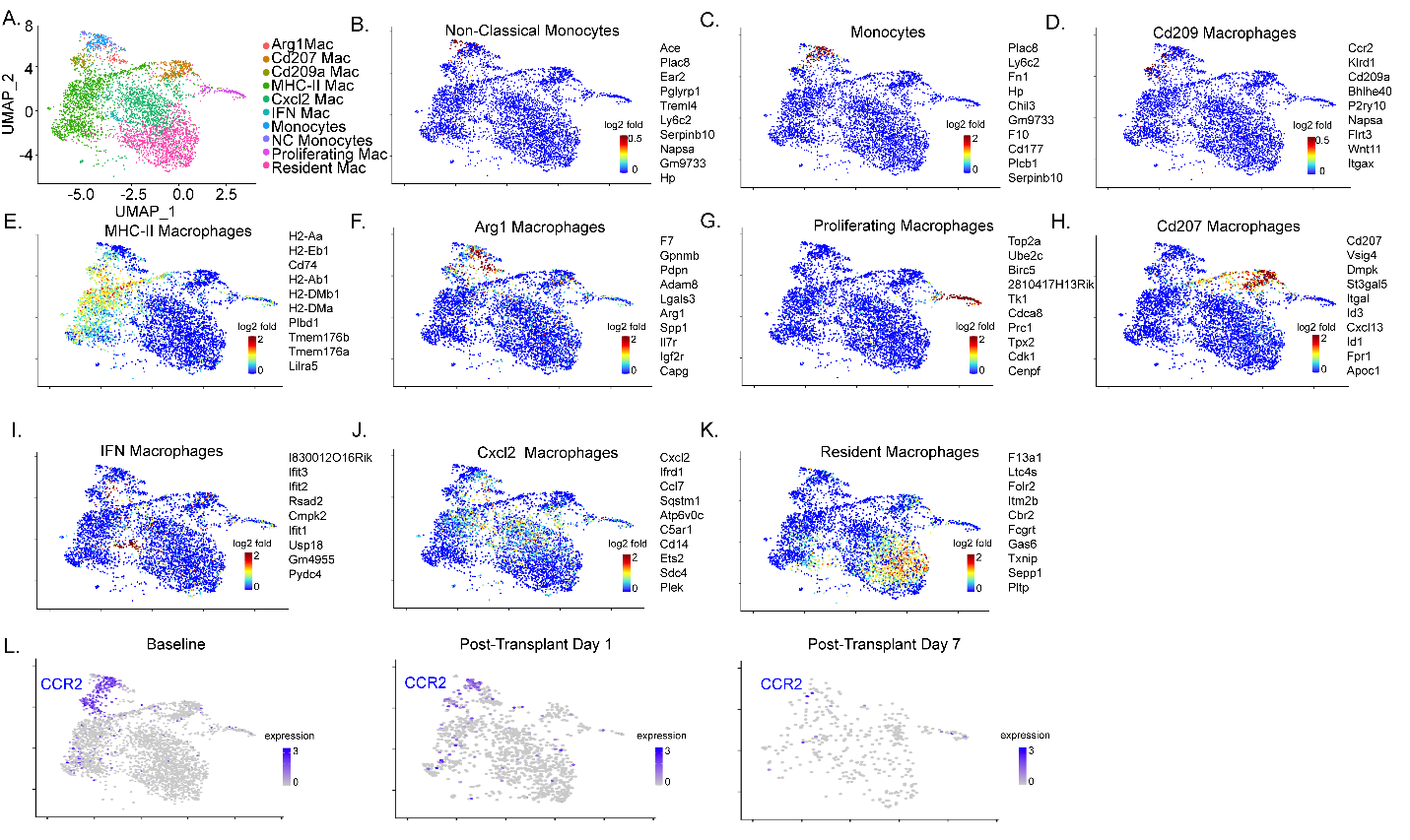

**Online Figure 7:** A) UMAP embedding plot for donors combined across all time points colored by cell type. Combined z-score for top 10 statistically significant differentially expressed genes overlaid on UMAP embedding plot in B) non-classical monocytes, C) classical monocytes, D) Cd209 macrophages, E) MHC-II^high^ macrophages, F) Arg1 macrophages, G) proliferating macrophages, H) Cd207 macrophages, I) IFN macrophages, J) Cxcl2 macrophages, and K) resident macrophages. L) UMAP embedding plot of Ccr2 expression at baseline, post-transplant day 1, and post-transplant day 7.

Online Figure 8:

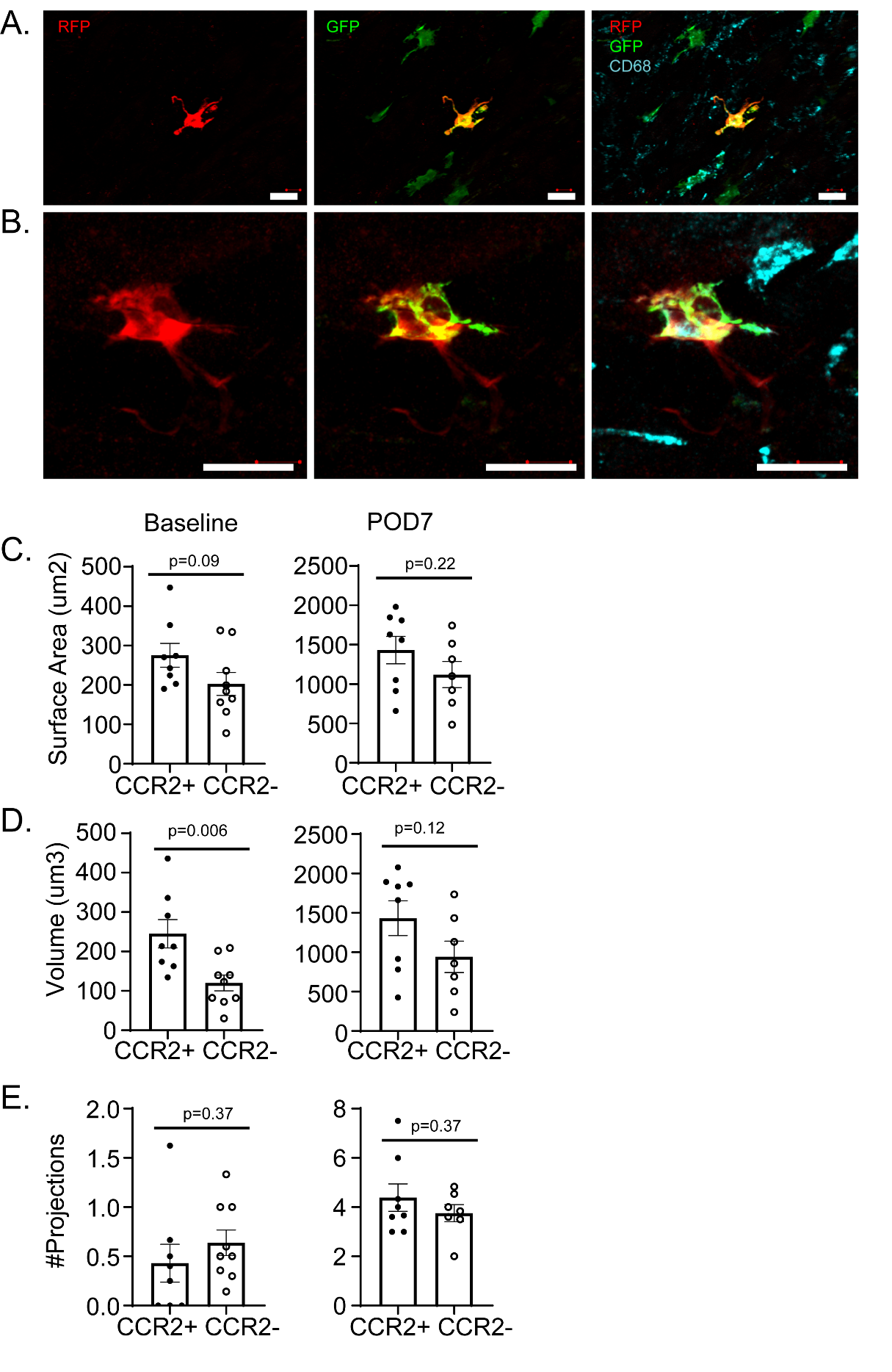

**Online Figure 8:** A) At post-transplant day 7, donor CCR2^-^ (GFP^+^ RFP^-^) macrophages surround a single donor CCR2^+^ macrophage (GFP^+^ RFP^+^). B) At post-transplant day 7, a donor CCR2^+^ macrophage is in proximity to recipient GFP^-^ RFP^-^ CD68^+^ macrophages. C) There are no significant differences in surface area measurements between donor CCR2^-^ macrophages and donor CCR2^+^ macrophages at baseline (Mann-Whitney U Test; p = 0.09,) or at d7 (Mann-Whitney U Test; p = 0.22). D) Donor CCR2^-^ macrophages have less volume than donor CCR2^+^ macrophages at baseline (Mann-Whitney U Test; p=0.006) but there are no differences in volume noted at d7 (Mann-Whitney U Test; p=0.12). E) There are no between group differences between donor CCR2^-^ and CCR2^+^ macrophages in terms of number of projections at baseline (Mann-Whitney U Test; p = 0.37) or at d7 (Mann-Whitney U Test; p = 0.37). Scale bar = 20 µm.

Online Figure 9:

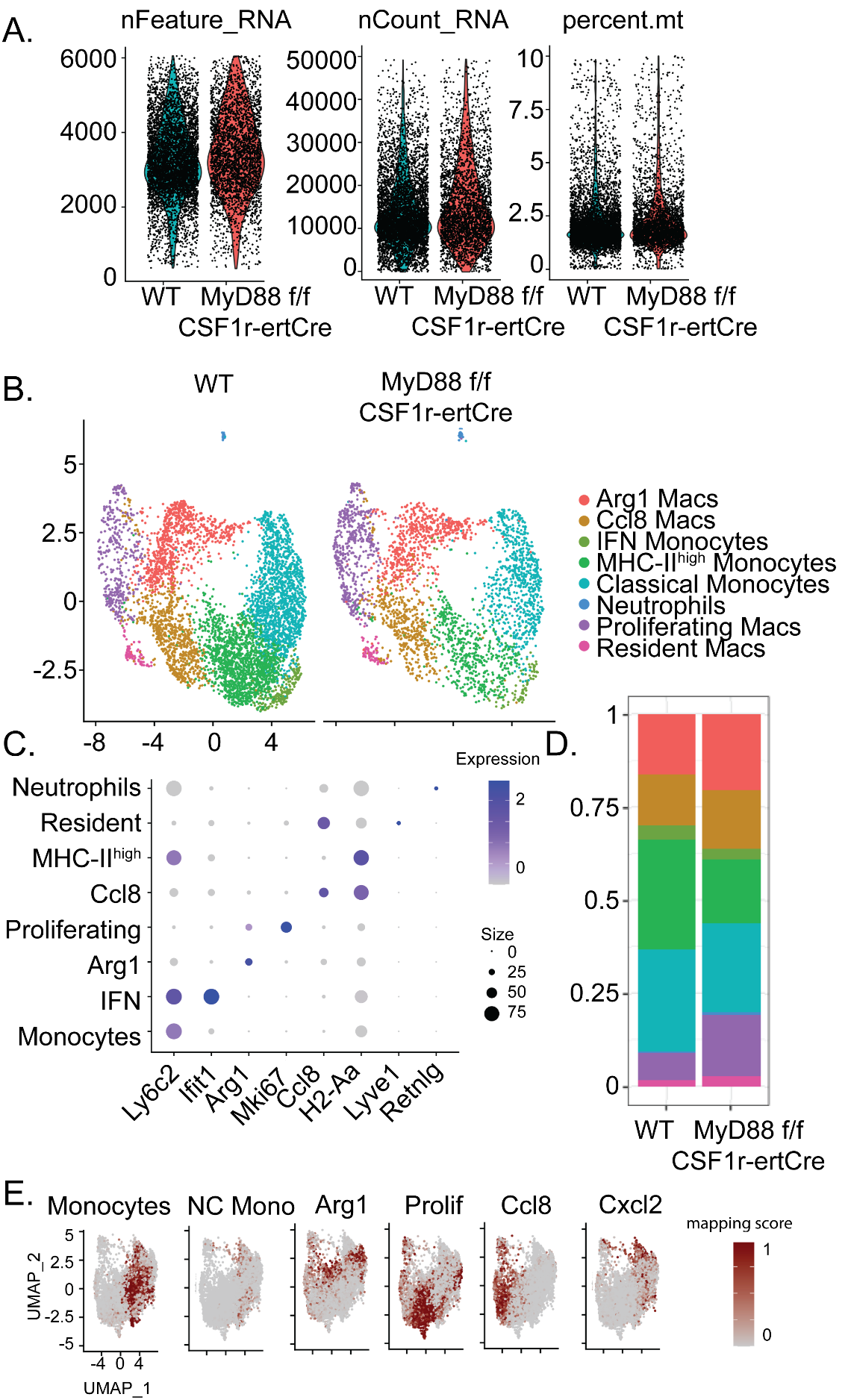

**Online Figure 9:** A) Quality control data plots showing feature counts, RNA counts, and percent mitochondria for each population. B) UMAP embedding plot of post-transplant day 3 WT and MyD88^f/f^ CSF1r^ertCre/+^ colored by cell type. C) Dot plot for cell type and top marker genes per cell cluster found from differential gene expression testing where dot size corresponds to percentage of cells expressing the gene and color corresponds to average expression in the given cluster. D) Bar plot of relative cell composition of each cluster by experimental condition. E) Mapping scores of post-transplant day 3 composite mapped onto recipient composite for monocytes, NC monocytes, Arg1 macrophages, proliferating macrophages, Ccl8 macrophages, and Cxcl2 macrophages.
